# Cytoneme-mediated signalling coordinates the development of glial cells and neurons in the *Drosophila* eye

**DOI:** 10.1101/2025.01.13.632741

**Authors:** Juan Manuel García-Arias, Gonzalo G. Girón, David Foronda, Isabel Guerrero, Antonio Baonza

**Affiliations:** Centro de Biología Molecular “Severo Ochoa”, CSIC /UAM, Madrid 28049, Spain; Departamento de Medicina, Facultad de Medicina Salud y Deportes, Universidad Europea de Madrid, Madrid 28670, Spain

**Keywords:** Cytonemes, Hh signalling, Glial cells, Eye imaginal disc, *Drosophila*

## Abstract

Effective cell communication is essential for the development and maintenance of the nervous system, where neurons and glial cells must interact closely. While cytoneme-mediated signalling is well-documented in various biological contexts, its role in coordinating neuron-glia development remains poorly understood. In this study, we investigated the function of cytonemes in neuron-glia coordination using the *Drosophila* eye imaginal disc as a model. This is a well-established system for examining the orchestrated development of glial and neuronal cells. Our results reveal that glial cells produce two distinct types of cytonemes based on their spatial orientation: one set extends toward nascent photoreceptors, while the other targets the morphogenetic furrow (MF). We have characterised the dynamics of glial cytonemes and demonstrated that disrupting these structures has a significant impact on glial cell migration and differentiation. This highlights the critical role of cytoneme-mediated signalling in regulating glial behaviour. Our findings also demonstrate that cytoneme function is essential for activating the Hedgehog (Hh) pathway in glial cells, with Hh ligand produced by photoreceptors. This pathway is necessary for glial differentiation, uncovering a previously unrecognised role for Hh signalling in this process. Overall, our results suggest that cytoneme-mediated Hh signalling is key to coordinating the development of both glial and neuronal populations.

## INTRODUCTION

Effective cell communication is essential for development and tissue homeostasis, ensuring proper organ size, shape and function. Precise intercellular signalling allows specific cells to send molecular signals that trigger targeted responses in receiving cells. This regulation is particularly important in the development of the nervous system. Within this complex neural network, neurons and glial cells work in synergy: neurons transmit and process information, while glial cells provide essential support, including maintaining homeostasis^1^. The formation of a functional nervous system depends on the coordinated interaction between these cell types. Unlike neurons, which remain localised, glial cells can migrate significant distances to reach their target locations, guided by a balance of attractive and repellent cues, often directed by neurons. As a result, the development of both glial cells and neurons becomes intricately intertwined and coordinated.

The *Drosophila* eye imaginal disc, the precursor of the adult eye^2^, offers a robust platform for investigating the mechanisms and signalling pathways that underpin neuronal and glial development. A significant body of research has demonstrated its value in advancing our understanding of glial biology in mammals^3–6^.

The eye disc arises from a subset of ectodermal cells that are uniquely specified during embryonic development, clearly distinguished from the neuroectodermal cells responsible for the formation of the central nervous system (CNS)^2^. Due to its developmental characteristics and origin, the eye disc can be considered as part of the peripheral nervous system (PNS), unlike the vertebrate eye, which is a component of CNS^7,8^. However, much like their mammalian counterparts, the eye disc contains both neurons and glial cells.

The process of neurogenesis begins in the third larval stage, initiated by the morphogenetic furrow (MF), an apical indentation moving from posterior to anterior. The MF triggers cell differentiation into neurons in a sequential manner^9^. Posterior to the MF, photoreceptor neurons and accessory cells organise into ommatidia, the structural units of the compound eye (reviewed in^10^). Photoreceptor axons extend through the optic stalk to form synapses across distinct regions of the optic lobe – the lamina and the medulla^11–14^.

The glial cells in the eye disc have their origin in the optic stalk, which connects the developing disc to the brain^15^. During the larval stages, these cells migrate to the eye disc as neurogenesis initiates. Once photoreceptor differentiation commences in the region behind the MF, glial cells leave the optic stalk and migrate onto the disc^7,14–17^.

The eye disc contains three main glial cell types, along with additional morphologically distinct varieties^16^. Two large subperineurial “carpet cells” cover the differentiated region, while perineurial glial (PG) cells, the migratory population, lie beneath them. As PG cells enter the disc, they move anteriorly to contact nascent photoreceptors (Fig. 1). This interaction triggers their differentiation into wrapping glia (WG), which adopt a morphology to encase photoreceptor axons, guiding them toward the optic ganglia in the CNS^5, 7,12,18^.

**Figure 1.**
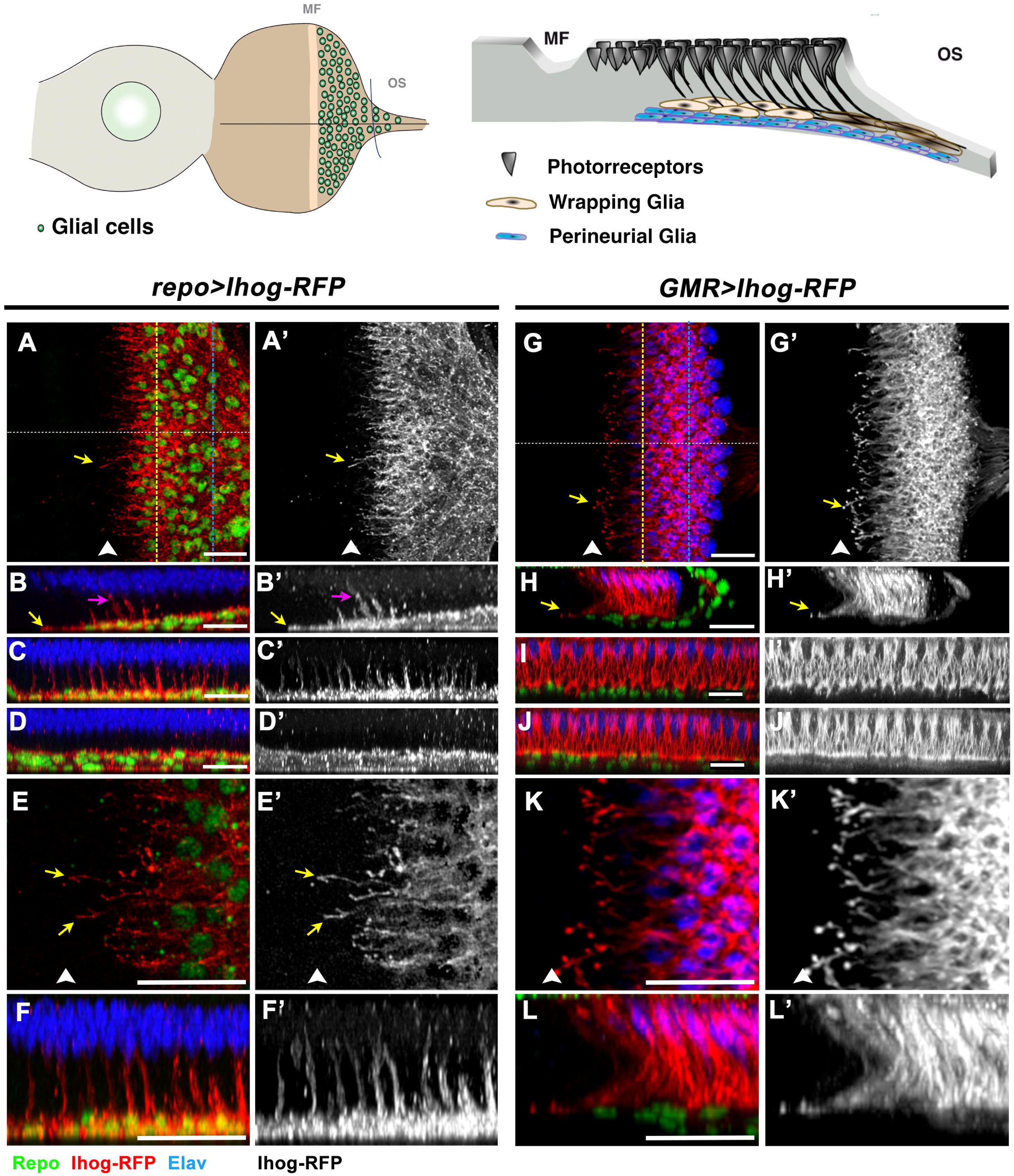
Glial and retinal cells form cytonemes in the Drosophila eye disc. The schematic illustration on the left represents an eye imaginal disc, with highlighted locations of the morphogenetic furrow and glial cells. The illustration on the right presents a transverse section, showing photoreceptors in the apical region and two main glial cell types in the basal region. (A, A’) Basal region of a *repo>Ihog-RFP* eye disc showing Repo (green), Ihog-RFP (red), and Elav (blue) staining. Glial cells extend cytonemes (yellow arrow) toward the morphogenetic furrow (MF, white arrowhead). (B, B’) Orthogonal view of the disc shown in A (white dashed line in A), with the optic stalk to the right. Glial cytonemes are directed toward the MF (yellow arrow) and photoreceptor cells (purple arrow). (C, C’) Orthogonal view of the disc shown in A (yellow dashed line in A). Glial cells at the region close to the MF produce photoreceptor-oriented cytonemes. (D, D’) Orthogonal view of the disc shown in A (blue dashed line in A). Photoreceptor-oriented cytonemes are absent in glial cells located in the posterior region of the eye disc. (E, E’) High magnification of A, A’ highlighting MF-oriented cytonemes (yellow arrows). (F, F’) Magnification of C, C’ showing photoreceptor-oriented cytonemes. (G, G’) *GMR>Ihog-RFP* eye disc showing Repo (green), Ihog-RFP (red), and Elav (blue) staining. Retinal cells extend cytonemes (yellow arrow) toward the MF (white arrowhead). (H, H’) Orthogonal view of the disc shown in G (white dashed line in G), optic stalk to the right. Retinal cells produce MF-oriented cytonemes (yellow arrow) in the basal region. (I, I’) Orthogonal view of the disc shown in G (yellow dashed line in G). (J, J’) Orthogonal view of the disc shown in G (blue dashed line in G). No glial-oriented cytonemes are observed. (K, K’) High magnification of G, G’. (L, L’) High magnification of H, H’. Both show MF-oriented retinal cytonemes. Scale bars: 20 μm.

These developmental features make the eye disc an outstanding model system for examining the intricate interplay between glial and neuronal cells, as well as the mechanisms responsible for coordinating their development.

The majority of signalling pathways that coordinate neuron and glial development in the eye disc rely on secreted ligands. These ligands regulate key processes such as proliferation, apoptosis, and migration by acting across specific distances and times, necessitating precise timing, intensity, and directionality. Gaining insight into how these molecules efficiently reach their target cells is crucial for unravelling the mechanisms underlying neuronal and glial coupling.

The diffusible factors Hedgehog (Hh) and the BMP2/4 morphogen Decapentaplegic (Dpp) play a pivotal role in regulating glial migration and proliferation^12,19–21^. Once differentiated, photoreceptors release Hh, which then spreads to influence cells anterior to the MF and glial cells^19,20,22,23^. Hh also induces Dpp expression ahead of the MF^24–27^. Collectively, Hh and Dpp enhance glial motility, with Dpp promoting proliferation^21,23^. It is notable that Hh signalling restrains glial migration prior to neurogenesis^20^, highlighting the integral role of these pathways in coordinating neuronal and glial development in the *Drosophila* visual system. However, the mechanisms that coordinate and regulate the precise timing, intensity, and directionality of these signalling pathways remain largely unexplored, despite their critical importance in determining the diverse outcomes they mediate.

Cytonemes are thin, specialized cellular projections essential for cell-to-cell signalling and communication. These structures are produced by both signal-emitting and -receiving cells, extending between them to enable direct contact at specific sites for signal reception and transduction^28^ (reviewed in ^29–32^). In the developing *Drosophila* wing disc, cytonemes have been implicated in mediating Hh, Dpp/BMP, Bnl/FGF, Spitz/EGF, Wg/WNT, and Notch signaling. Additionally, there is growing evidence for cytoneme involvement in morphogen transport in vertebrates (reviewed in ^29^). Despite their crucial role in regulating intercellular signaling across various tissues, their presence and function in nervous system development remain largely unexplored. Notably, the role of cytoneme-mediated signaling in neuron-glia interactions during development is unknown. The distinctive developmental features of the eye disc make it an excellent model for investigating whether cytonemes coordinate signals that integrate neuronal and glial development.

In this study, we used the eye disc to explore cytonemes involvement in the signalling pathways that regulate neuron-glia communication, coordinating their differentiation and migration. Our findings indicate that glial cells produce two distinct sets of cytonemes, distinguished by their spatial orientation. The first set of cytonemes, directed towards the MF, is likely to facilitate the reception of signals that are crucial for glial positioning and migration. The second set, oriented towards nascent photoreceptors, activates the Hh pathway in glial cells through Hh ligand produced by photoreceptors. This activation promotes the differentiation of perineurial glial cells into wrapping glia (WG). These results reveal a previously unrecognised function of Hh signalling in regulating glial differentiation. Through cytoneme-mediated signalling, these specialised glial cytonemes coordinate both migration and differentiation – two critical behaviours essential for the proper development of glial.

## RESULTS

### Both glial and retinal cells produce cytonemes in the *Drosophila* eye disc

To investigate the potential role of cytoneme-mediated signalling in neuron-glia interactions, we first examined the presence of cytonemes in both glial and retinal cells in the eye disc. This was achieved by expressing a fluorescently tagged form of the Hh co-receptor protein Ihog (Ihog-RFP), known to stabilised cytonemes in wing discs and abdominal histoblasts^33^, in either glial and retinal cells using the *repo-Gal4* and *GMR-Gal4* lines, respectively. We observed that glial cells, identified by positive Repo staining, produced two distinct types of cytonemes with different spatial orientations: a set of cytonemes oriented towards the MF (MF-oriented cytonemes) (yellow arrows in Fig. 1A-A’ and B-B’, and Video 1), localised in the basal region of the disc, and a second set that projects vertically towards the photoreceptor neurons (PR-oriented cytonemes), identified by positive Elav staining (purple arrows in Fig. 1B-B’). Interestingly, these later cytonemes are produced only by the most anterior glial cells, closer to the MF, whereas posterior glia cells do not produce them (Fig.1B-B’ and compare anterior section Fig. 1C-C’ with posterior section Fig. 1D-D’ and Video 1).

MF-oriented cytonemes form a dense array with an average maximum length of approximately 22,3±3,1 μm (Std.Dev, n=21 discs). Some cytonemes can extend to lengths of up to 30 μm. The average length of PR-oriented cytonemes is 17,9±6,5 μm (Std.Dev, n=19 discs), although some can be longer up to 27 μm (Fig. S1). Interestingly, we observed that some PR-oriented cytonemes are branched, especially towards their distal end (Fig. 1F-F’).

We also observe that retinal cells generate a dense array of cytonemes that project through the basal region of the disc (Fig. 1G-L’). These cytonemes always extend toward the MF (yellow arrows in Fig. 1G-H’) with an average maximum length of approximately 14,4+1,6 μm (Std.Dev, n=17 discs). The tips of these cytonemes are often thickened, forming a sphere (Fig. 1K-L’). Intriguingly, our observations indicate the absence of cytonemes directed towards glial cells (Fig. 1I-L’).

As previously described^34^, fluorescently tagged Ihog labelled punctate structures, some of which were found in cytonemes, while others were not associated with them (Fig. 1A-F’). These structures may correspond to exovesicles, as previously described^34^.

To determine if these filopodial structures are actin-based, as indicated by previous studies^31,33^, we expressed the actin-binding markers *UAS-Moe-Cherry* and *UAS-Lifeact-mRFPruby*^35^ in glial and retinal cells using *repo-Gal4* and *GMR-Gal4* lines. The distribution of cytonemes labeled with these markers closely matches that of *Ihog-RFP*-labeled cytonemes in both glial cells (Fig. 1, 2, and Fig. S2A-F’) and retinal cells (Fig. S2G-L’), confirming their actin-based nature. This also indicates that these structures are not artefacts of Ihog overexpression. Furthermore, Ihog-RFP overexpression had no discernible impact on glial cell number or migration, regardless of whether it was expressed under the *repo-Gal4* or *GMR-Gal4* lines (Fig. S1).

**Figure 2.**
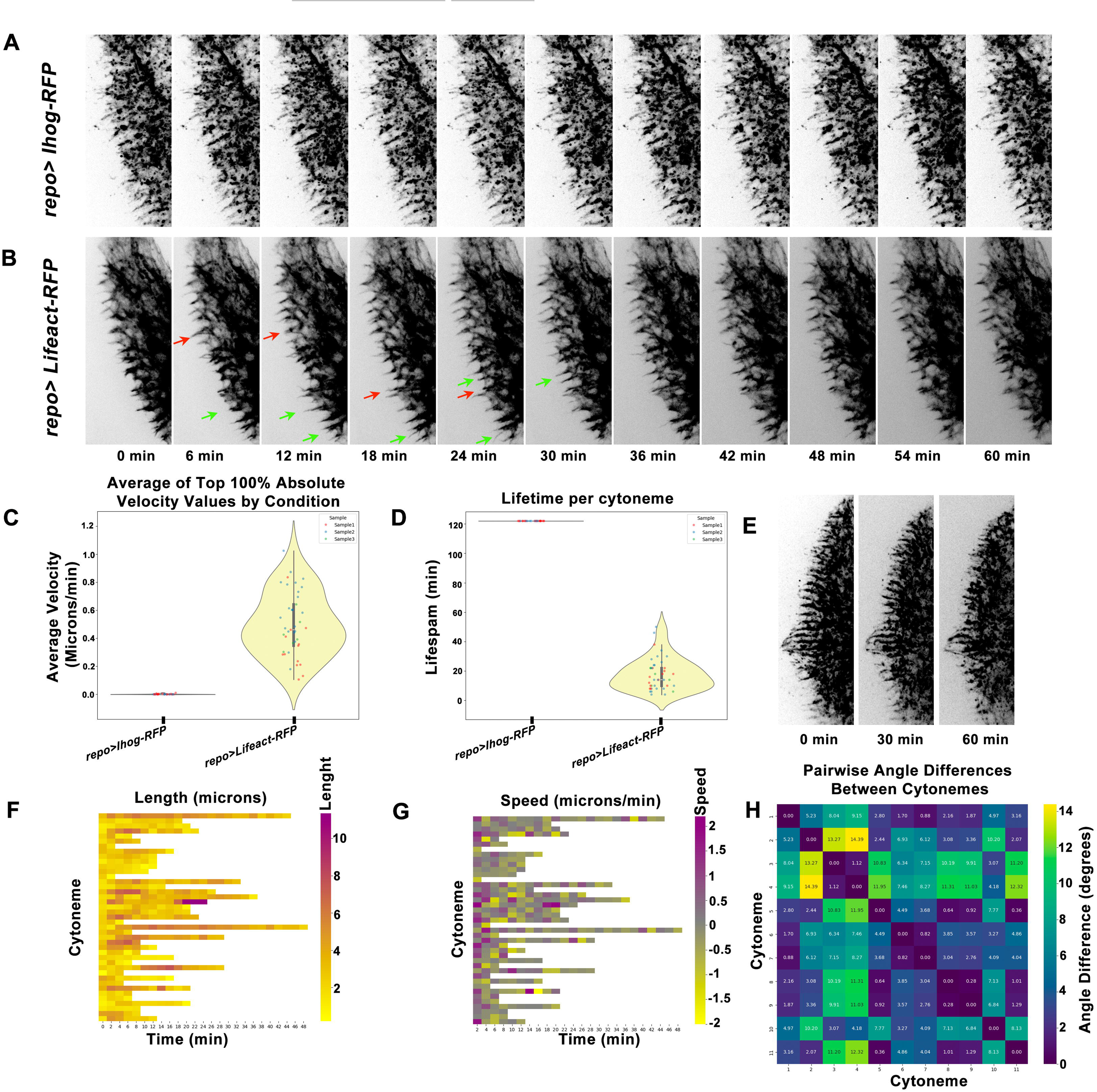
Dynamic Behavior of Glial Cell-Produced Cytonemes. (A-B) Representative frames from Supplementary Videos 2 (A) and 3 (B) show *ex vivo* imaging of L3 stage eye discs cultured at 25°C for 60 minutes. (A) Cytonemes labeled with *repo-Gal4>UAS-Ihog-RFP* are highly stable, showing no structural or length changes during the observation period. (B) Cytonemes labeled with r*epo-Gal4>UAS-Lifeact-mRFPruby* exhibit dynamic behavior, continuously elongating (green arrows) or retracting (red arrows). Length changes occur rapidly, with some cytonemes altering noticeably within 6 minutes. (C-D) Violin plots comparing the average velocity (C) and lifespan (D) of cytonemes labeled with Ihog-RFP and Lifeact-mRFPruby (n=30). (E) Time-lapse frames showing parallel alignment of cytonemes labeled with Ihog-RFP, maintaining orientation over time. (F) Length heatmap: x-axis indicates time, y-axis represents individual cytonemes, and color maps cytoneme lengths (yellow: up to 12 μm, purple: close to 0μm). (G) Velocity heatmap: x-axis represents time, y-axis corresponds to individual cytonemes, and colors depict velocity (purple: elongation up to 2 μm, grey: minimal movement, yellow: retraction up to −2 μm). (H) Angle heatmap: x-axis represents time, y-axis shows cytonemes in (E) at Time 0 min. Colors indicate angle differences (purple: near 0°, almost parallel; yellow: up to 14°).

To study the dynamic behavior of cytonemes emanating from glial cells, we observed them over time using live imaging (Video 2 and 3). We monitored the actin cytoskeleton by utilizing fluorescently labeled *UAS-Ihog-RFP* or *UAS-Lifeact-mRFPruby*, expressed under the control of the *repo-Gal4*. We then analysed the behaviour of individual cytonemes. Using the *Ihog-RFP* construct, we observed single protrusions that remained very stable for up to 2 hours without growing or shrinking (Fig. 2A and C-D, and Video 2). This is consistent with previous works in wing discs and abdomen, in which Ihog-RFP expression stabilised cytonemes in glial cells^34,33^. Therefore, to study cytoneme dynamics more effectively, we used *UAS-Lifeact-mRFPruby*^35^ instead of *Ihog-RFP*. We found that glial cytonemes tagged with *Lifeact-mRFPruby* are highly dynamic, with a lifetime of approximately 17.76 ± 11 minutes (n=30) (Fig. 2B-D and Video 3). Over time, these cytonemes continuously grow and shrink (Fig. 2F and G, Video 3) until they reach their final length. The growth rate of these cytonemes is approximately 0.49±0.22 μm/min (Fig. 2G).

We took advantage of the high stability of the cytonemes, achieved by expressing *Ihog* in glial cells, to study their distribution over time. We observed that these cytonemes maintain a parallel distribution for extended periods of time, up to 2 hours Fig. 2 E and H, and Video 2). This consistent parallel arrangement suggests the existence of a regulated mechanism that may be involved in producing cytonemes in an organized pattern as it has been described at the A/P compartment border of the wing imaginal disc^36^.

### Glial cytonemes play a role in regulating both the number and positioning of glial cells

To investigate the role of cytonemes in glial development, we first inhibited the formation of cytonemes produced by glial cells and then analysed glial behaviour. To this end, we used *repo-Gal4* to express different *UAS-RNAi* constructs in glial, targeting Rack1 (*Rack1^RNAi^*), Coracle (*cora^RNAi^*) and Pico/Lamelipodin (*Pico^RNAi^*), all of which have previously been reported to affect cytoneme homeostasis^29,34,37^. To minimise potential side effects, we used the *tub-Gal80^ts^* factor to transiently restrict the expression of these RNAi constructs to a 24-hour timeframe. We observed that glial cytonemes were affected in all the genetic conditions analysed (Fig. S3). Specifically, depleting *cora* or *Pico* (*repo^ts^-Gal4>UAS-Ihog-RFP UAS-cora^RNAi^* and *repo^ts^-Gal4>UAS-Ihog-RFP UAS-Pico^RNAi^*) significantly reduced the maximum length of MF-oriented cytonemes and the number of PR-oriented cytonemes (Fig. S3C-D’’’ and E-F). *Cora* depletion also decreased the number of MF-oriented cytonemes, while *Pico* had no effect on them (Fig. S3F-G). Down-regulating (*Rack1 repo^ts^-Gal4>UAS-Ihog-RFP UAS-Rack1^RNAi^*) reduced the number of both MF- and PR-oriented cytonemes but did not affect their length (Fig. S3B-B’’’ and E-F).

We then analysed the number and position of glial cells in the discs of the genotypes described above. We found a significant reduction in the number of glial cells in all conditions examined (Fig. 3 A-E). This reduction is not due to an increase in apoptosis or a decrease in cell proliferation, as evidenced by the absence of dead glial cells, as assessed by staining for the anti-cleaved effector caspase Dcp1 (Fig. S4 A-C’ and D), and unaffected glial proliferation, as shown by staining for the mitotic marker PH3 (Fig. S4 E-H). These findings suggest that a defect in glial migration or glial motility is more likely responsible for the reduction in glial cell numbers. Accordingly, we found that in these discs the glial cells fail to migrate to their normal position within the eye disc. Whereas in *repo-Gal4>UAS-Ihog-RFP* control discs most anterior glial cells remain 3 or 4 photoreceptor rows behind the MF (white dashed line in Fig. 3 A’ and F), glial cells in *repo^ts^-Gal4>UAS-Ihog-RFP UAS-Rack1^RNAi^* and *repo^ts^-Gal4>UAS-Ihog-RFP UAS-Pico^RNAi^* remain a significant number of rows behind this position (white dashed lines in Fig. 3 B’-C’ and F).

**Figure 3.**
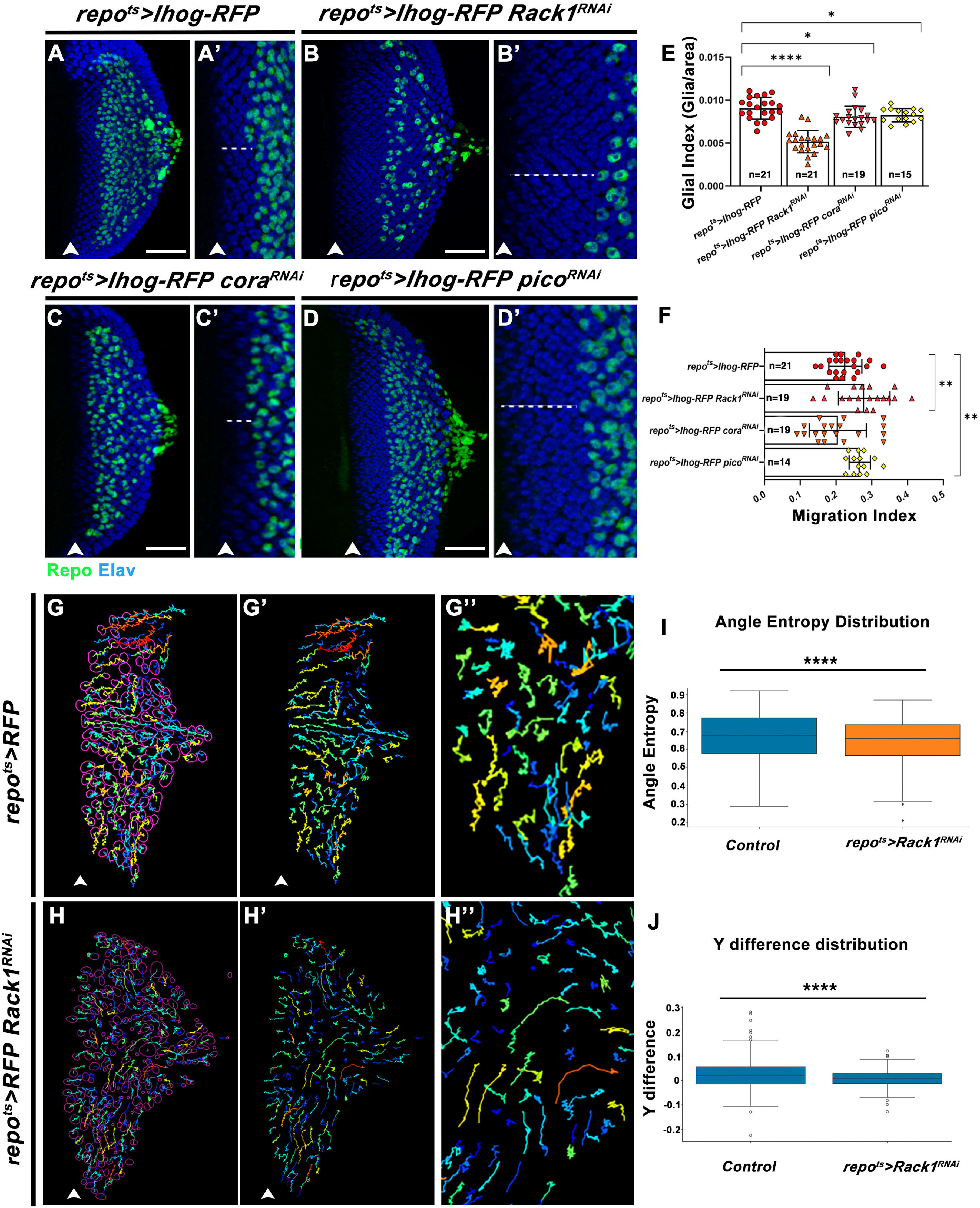
Inhibition of glial cytonemes reduces glial numbers and impairs migration. (A-D’) Eye imaginal discs of *repo>Ihog-RFP* (A, A’), *repo>Ihog-RFP Rack1^RNAi^* (B, B’), *repo>Ihog-RFP cora^RNAi^* (C, C’), and *repo>Ihog-RFP pico^RNAi^* (D, D’) genotypes, showing Repo (green) and Elav (blue) staining. The MF is on the left (white arrowhead). Dashed lines in A’-D’ indicate the distance between the most anterior glial cell and the first photoreceptor row. Scale bars: 50 µm. (E-F) Quantification of glial index (E) and glial migration index (F) for the indicated genotypes. Statistical significance: *p*<0.05; **p**<0.01; \*\***p**>0.0001. (G-H’’) Tracking data of RFP-labeled glial cells from *repo-RFP* (G-G’, Video 4) and *repo-RFP Rack1^RNAi^* (H-H’, Video 6) discs. Discs were imaged *ex vivo* at L3 stage for 90 minutes (Video 5 and Video 6). Purple circles mark glial positions at *t*=0. In G’ and H’, only cell trajectories are shown. G’’ and H’’ display magnified views of cell movement trajectories. (I-J) Graphs quantifying glial movement. (I) Angle entropy distribution shows averages of 0.67 (control) and 0.64 (*repo^ts^>Gal4 UAS-Rack1*^RNAi^), ***p***=0.0002. (J) Displacement towards the MF is expressed as a ratio of glial movement distance to the total distance from the optic nerve to the MF. Averages: 0.029 (control) and 0.0079 (*repo^ts^>Gal4 UAS-Rack1*^RNAi^), ***p***=0.0007.

Taken together, these findings suggest that glial cytonemes play a critical role in regulating the glial cell numbers during eye disc development.

### Retinal cytonemes do not have an impact on glial development

Next, we investigated whether retinal cytonemes play a role in determining glial cell number and/or migration behaviour. To test this, we disrupted cytoneme formation by expressing the *UAS-Rack1^RNAi^* construct under the control of the *GMR-Gal4* driver, selected for its specific expression in retinal cells. To minimise potential side effects, we used the *tub>Gal80^ts^*factor to restrict the expression of the construct to a 24-hour window.

As expected, eye discs of this genotype showed a significant reduction in Ihog stabilised cytonemes, both in length and index (Fig. S5 A-D), consistent with our previous findings using the *repo-Gal4* driver. Despite this reduction, the glial index remained unaffected by retinal cytoneme inhibition (Fig. S6 A-C). Notably, glial cells in *GMR^ts^>UAS-Ihog-RFP UAS-Rack1^RNAi^*discs showed a slight but statistically significant tendency to migrate to more anterior positions within the eye disc compared to control *GMR^ts^>UAS-Ihog-RFP* discs (white dashed lines in Fig. S6 A’’, B’’ and D).

These results suggest that inhibition of retinal cytonemes has a minimal impact on glial development and does not affect the number of glial cells. However, given their orientation towards the MF, it is likely that they may contribute to its progression.

### Disruption of cytonemes affects the migratory behaviour of glial cells

We monitored the trajectory and migration behaviour of both control glial cells and those with altered cytoneme formation by live imaging during a 90-minute session of *ex vivo* cultured third-instar eye discs (Video 4-7 and Fig. 3 G-H’’). To analyse their migration patterns, we calculated the entropy of their angular distribution - an approach that integrates cellular turning dynamics with Shannon entropy. This approach allows us to study the statistical and temporal properties of glial migration persistence in a complex environment^38^. We observed that the angular entropy distribution is higher in control glial cells compared to those with defective cytonemes (Fig. 3 I). This suggests that the movement directions of control glial cells are more random or less coordinated than in the mutants. We observed that while glial cells with defective cytonemes exhibit relatively straight and regular trajectories, control glial cells exhibit highly zigzag-like trajectories with frequent changes in direction (Fig. 3 compare G’’ with H’’). Although this may seem counterintuitive, since defective cytonemes are expected to impair the regulation of cell migration via intercellular signals, one possible explanation is that control glial cells receive strong but conflicting signals from multiple sources, leading to higher angular entropy. In contrast, glial cells with defective cytonemes may experience reduced intercellular regulation, thereby reducing the complexity of the signals they receive and resulting in a lower angular entropy distribution.

We also analysed the mobility of glial cells towards the MF, focusing exclusively on those adjacent to the optic nerve. To account for variations in the size of the eye discs analysed, we normalised the data by defining the Y difference as the ratio between the distance that glial cells moved towards the MF and the distance between the optic nerve and the MF (total movement in μm/ distance from the optic nerve to the MF in μm). Our results showed that control glial cells moved slightly, but statistically significantly, more anteriorly than glial cells with impaired cytonemes (Fig. 3 J).

Overall, our results suggest that the migratory behaviour of glial cells with disrupted cytonemes is altered, being less complex and having less motility towards the MF.

### Glial cytonemes are needed for proper perineural glial differentiation into wrapping glia in the eye disc

As neuronal development begins, some perineurial glial cells located in the most anterior region undergo a process of differentiation, changing from a migrating cell type to a differentiating wrapping glial cell type ^6,7^. The differentiation of this glial cell type has been suggested to depend on the physical interaction between perineurial glia and nascent photoreceptor axons^6,7^. Given our identification of a subset of cytonemes directed towards photoreceptor cells, we investigated whether these glial cytonemes might be involved in this process.

To this end, we analised the cytonemes produces by the two main glial cells located in the eye discs, the PG and WG glial cells. We used the specific *Mz97-Gal4* and *c-527-Gal4* lines to overexpress Ihog-CFP in WG and PG, respectively^14,16^. Our analysis showed that WG cells do indeed produce cytonemes, with a spatial distribution similar to that of the entire glial population (Fig. 4 A-D’). We observed both MF- and PR-directed cytonemes; interestingly, MF-directed cytonemes from WG cells appear slightly shorter compared to those from the broader glial population, as their average maximum length is 14,6±1,9 μm (Std.Dev, n=20 discs), (compare Fig. 4 A-D’ with Fig.1 A-C). We observed that PG also displayed both MF- and PR-oriented cytonemes (Fig. 4 E-H’ and I for the illustrative scheme).

**Figure 4.**
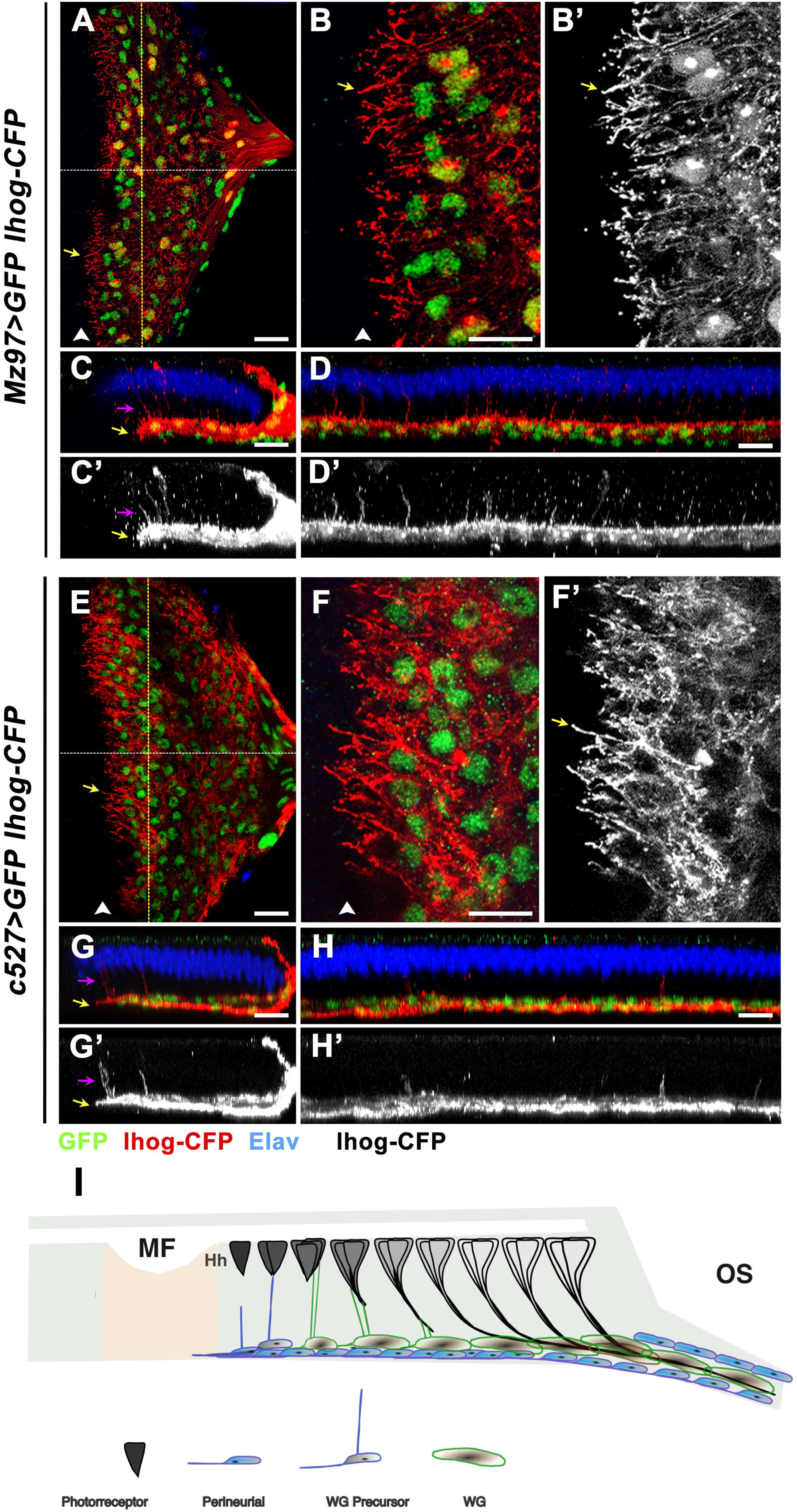
Cytonemes produced by perineural and wrapping glial cells. (A-B’) Basal region of an *Mz97>GFP Ihog-CFP* eye disc showing GFP (green), Ihog-CFP (red), and Elav (blue). Wrapping glial cells (WG) produce cytonemes (yellow arrows) directed toward the MF (white arrowhead). (B, B’) Magnified views of (A). Scale bars: 20 µm (C-D’) Orthogonal views of the disc shown in A. (C, C’) White dashed line in (A), optic stalk to the right. WG cytonemes target both the MF (yellow arrows) and photoreceptor cells (magenta arrows); photoreceptor-oriented cytonemes are only produced by the most anterior WG. (D, D’) Orthogonal views of the disc shown in A (Yellow dashed line). Scale bars: 20 µm (E-H’) Basal region of a *c527>GFP Ihog-CFP* eye disc showing GFP (green), Ihog-CFP (red), and Elav (blue). (E-F’) Perineural glial cells (PG) produce MF-oriented cytonemes (yellow arrows, white arrowhead). (F, F’) Magnified views of (E). (G-H’) Orthogonal views of the disc shown in E. (G-G’) White dashed line in E. (H-H’) Orthogonal views of the disc shown in E (yellow dashed lines). PG produce fewer photoreceptor-oriented cytonemes (magenta arrows). Yellow arrows indicate MF-oriented cytonemes. Scale bars: 20 µm (I) Schematic representation showing cytoneme production by WG and PG. Both cell types produce cytonemes targeting the MF and photoreceptors. The optic stalk (OS) is to the right.

Altogether, our results suggest that both PG and WG can produce MF- and PR-directed cytonemes (Fig. 4 I).

To assess the impact of the cytonemes generated by WG on their differentiation, we suppressed their formation by overexpressing *Rack1^RNAi^*under the control of the *Mz97-Gal4* line. Interestingly, we observed a significant decrease in the number of WG cells upon *Rack1* depletion (*Mz97-Gal4>UAS-GFP UAS-Rack1^RNAi^*), while the total number of glial cells remained unaffected (Fig. 5 A-D). As a result, the proportion of WGs over the total glial cell number was greatly reduced (Fig. 5 E). These results suggest that the reduction in WG is due to an impaired differentiation process rather than reduced glial migration, thus implicating cytonemes in the differentiation of WG.

**Figure 5.**
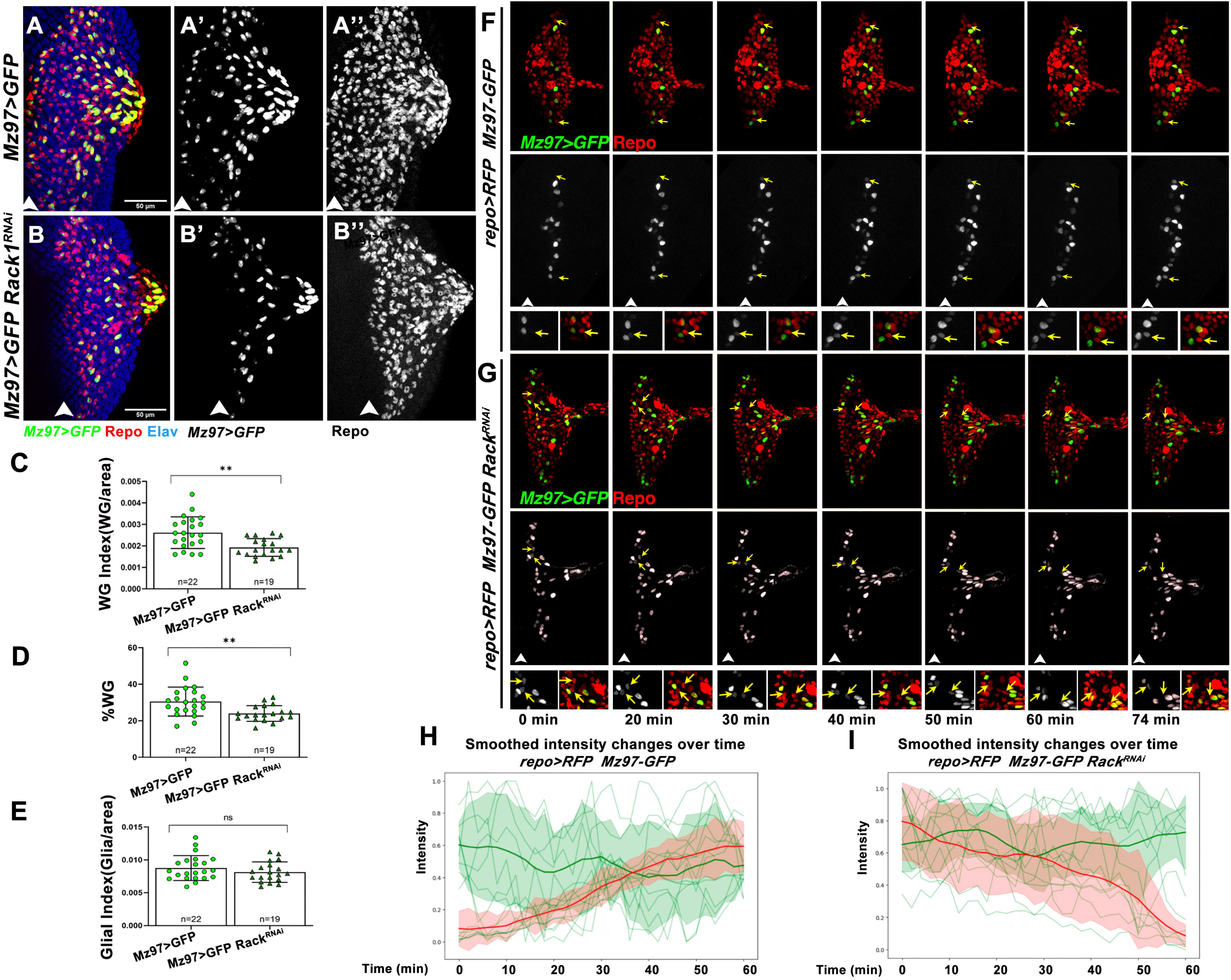
Interference with cytonemes formation in WG cells results in defects in glial differentiation. (A-A’’) *Mz97>GFP Ihog-RFP* eye discs showing GFP (green), Repo (red), and Elav (blue). MF is on the left (white arrowheads). (B-B’’) *Mz97>GFP Ihog-RFP Rack1^RNAi^* discs display reduced WG differentiation. (C-E) Quantifications of WG index (C), %WG (D), and glial index (E) for the genotypes described above. ns = not significant; **p<0.01. (F-G) Frames from Videos 8 and 9 showing *repo-RFP Mz97>GFP* (F, control, video 8) and *repo-RFP Mz97>GFP Rack1^RNAi^* (G, video 9) discs cultured *ex vivo* for 100 minutes. Glial cell nuclei are labelled with *repo-RFP* (red), while WG (wrapping glia) are labelled with *Mz97>GFP* (green). In control discs, some PG cells activate *Mz97>GFP*, differentiating into WG (yellow arrows, F). In *repo-RFP Mz97>GFP Rack1^RNAi^* discs, some glial cells show a decrease in *Mz97>GFP* expression over time (yellow arrows in G), indicating impaired differentiation. High-magnification insets highlight the changes. (H-I) Quantification of *Mz97>GFP* intensity over time. (H) In controls, two populations emerge: one with stable GFP expression (green, average intensity shown by the thick green line) and another with increasing GFP expression (red, thick red line). (I) In *repo-RFP Mz97>GFP Rack1^RNAi^* discs, a population retains stable GFP expression (green, thick green line), while another population exhibits declining GFP intensity over time (red, thick red line). Scale bars: 50 µm.

To further investigate this idea, we performed live imaging to follow the differentiation process of WG in both control and *Rack1*-depleted glial cells over time. We expressed *UAS-Rack1^RNA^*^i^ and *UAS-GFP* using the *Mz97-Gal4* line in *repo-RFP* flies, which express the RFP protein under the control of the *repo* promoter^39^ (Video 8 and Video 9). Consistent with our previous observations, we found that in control flies (*repo-RFP Mz97>GFP*), some glial cells gradually acquire GFP expression over time (Fig. 5 F and H, and Video 8). However, this process does not occur in glial cells depleted of *Rack1*. Furthermore, in some WG-expressing cells where low levels of GFP are initially observed, GFP expression decreases over time (Fig. 5G and I, and Video 9).

All together our results suggest that cytonemes play a key role in the process of differentiation of WG during eye development.

### Glial cytonemes may be implicated in Dpp and Hh signalling reception

Cytonemes have been shown to mediate the Hh and Dpp signalling pathways in several systems, including the wing disc^29,40^. Both pathways are critical for regulating glial cell number and migration during normal development and regeneration^21,23^. Based on this, we investigate whether glial cytonemes may play a role in regulating Hh and/or Dpp signalling during glial development.

First, we analysed the expression of Hh and Dpp ligands in the eye disc in relation to the spatial distribution of glial cytonemes. To analyse Hh expression we used a *BAC Hh-GFP* construct which mimics endogenous Hh distribution^41^. *Hh-GFP* forms an anterior-posterior gradient in the apical region of the eye disc, with higher expression in newborn photoreceptors compared to more mature ones (compare yellow and white arrows in Fig. 6 A’). This gradient is particularly pronounced in the apical region of the photoreceptors (compare Fig. 6 A’).

**Figure 6.**
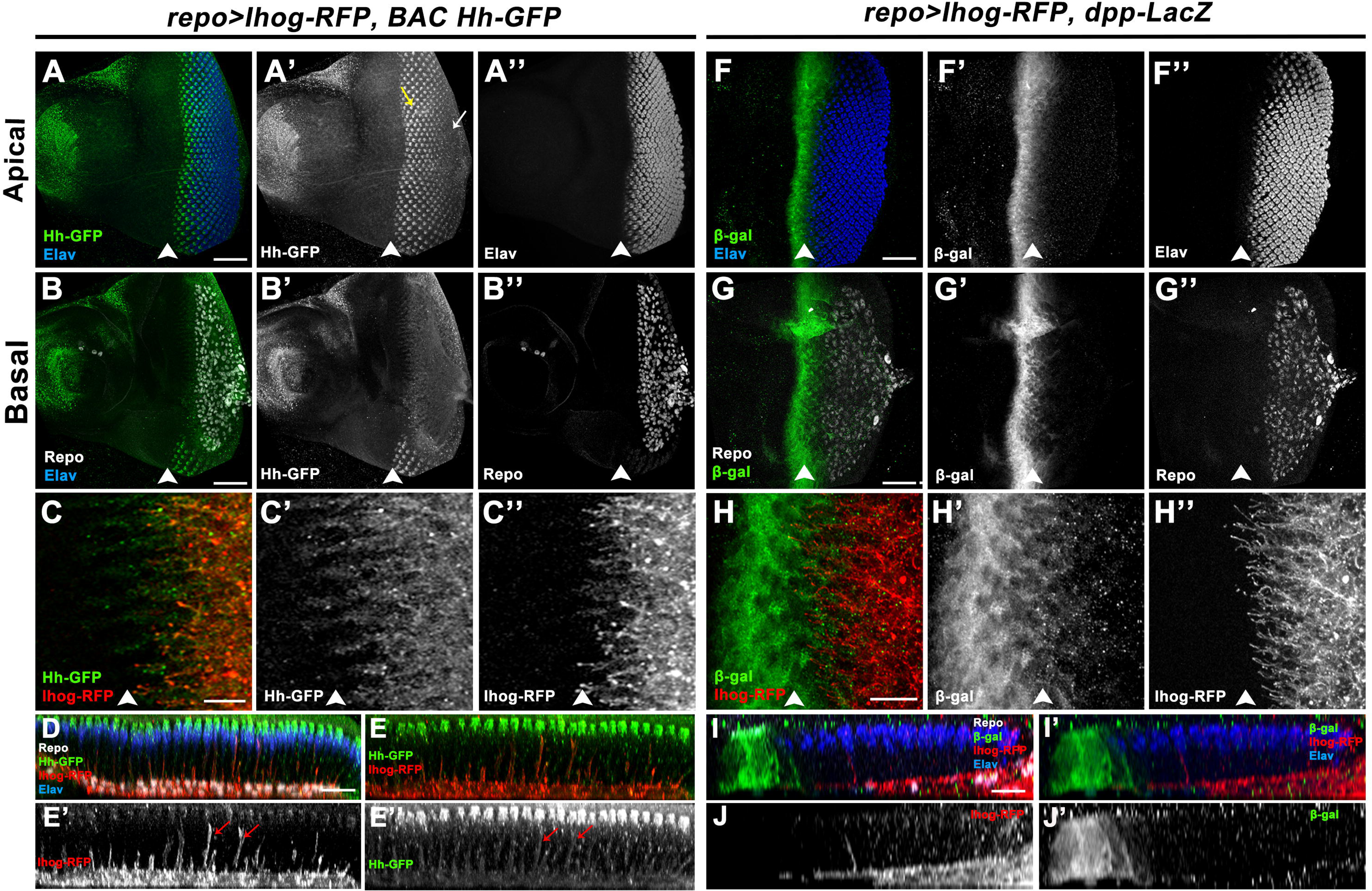
Glial cytonemes are oriented towards the sources of Dpp and Hh ligands. (A-E’’) *repo>Ihog-RFP BAC Hh-GFP* eye discs stained for Repo (gray), Hh-GFP (green), Ihog-RFP (red), and Elav (blue). The MF is positioned on the left (white arrowheads). (A-A’’) Hh-GFP forms a gradient in the apical region, decreasing from posterior (white arrow in A’) to anterior (yellow arrow in A’). (B-B’’) Hh-GFP expression in the basal region of the same disc. (C-C’’) MF-oriented glial cytonemes do not colocalize with Hh-GFP, which localizes to horizontal stranded structures extending towards the MF (compare C’with C’’). (D-E’’) Orthogonal views of the disc shown in A show photoreceptor-oriented cytonemes colocalizing with Hh-GFP along vertical stranded structures (red arrows, compare E’ with E’’). (F-J’’) *repo>Ihog-RFP dpp-LacZ* eye discs stained for Repo (gray), *dpp-LacZ* (green), Ihog-RFP (red), and Elav (blue). (F-F’’) Apical dpp-LacZ expression is observed in the cytoplasm. (G-G’’) Basal region of the same disc shows dpp-LacZ expression in the cytoplasm. (H-H’’) MF-oriented glial cytonemes connect with cells expressing dpp-LacZ. (I-J’’) Orthogonal views of the disc shown in F reveal MF-oriented glial cytonemes contacting dpp-LacZ-expressing cells. Scale bars: 50 µm (A-B’’, F-G’’) and 20 µm (C-E’’, H-J’’).

In the basal region of the disc, we observed that *Hh-GFP* is expressed in projections that extend towards the morphogenetic furrow, marked by punctate staining. These processes do not co-localise with the MF-oriented cytonemes produced by glial cells (Fig. 6 C-C’’), suggesting that these processes probably originate from retinal cells. However, in orthogonal sections, we observed that PR-oriented cytonemes of the glial cells co-localised with vertical *Hh-GFP* structures, and these cytonemes were able to reach the apical accumulations of Hh (Fig. 6 D-E’’).

To examine the expression of *dpp*, we used the *dpp-LacZ* reporter line, which is expressed in a band of cells anterior to the morphogenetic furrow and is strongly downregulated posterior to the furrow (Fig. 6 F-H’’). We observed that the MF-oriented cytonemes produced by glial cells extend to contact the band of *dpp*-expressing cells (Fig. 6 H, H’’). This is more clearly visualised in orthogonal sections of these discs (Fig. 6 I-J’).

These results suggest that the PR-oriented and MF-oriented cytonemes produced by glial cells are targeted to cells producing Hh and Dpp, respectively. Therefore, regarding PR-oriented cytonemes, it is plausible that they play a role in mediating Hh signalling during glial development. To investigate this further, we analysed whether inhibiting cytoneme formation affects Hh activation in glial cells. To this end, we examined the expression of the Hh receptor Patched (Ptc) and Cubitus Interruptus (Ci)^42–44^, two known targets of this pathway (reviewed in ^45^), in control and impaired glial cytoneme formation discs. In control discs (*repo^ts^-Gal4>UAS-GFP*), Ptc levels were significantly higher in glial cells close to the MF compared to those in more posterior regions (Fig. 7 A-A’; compare purple arrows with red arrows in Fig. 7 B’). Accordingly, when we calculated the ratio of Ptc levels between glial cells in the anterior region and those in the posterior region (A/P ratio), the average value is higher than 1 (Fig. 7 G). However, in *repo^ts^-Gal4>UAS-GFP UAS-Rack1^RNAi^* discs (Fig. 7 C-C’’’ and D-D’’), Ptc expression in glial cells adjacent to the MF was reduced compared to control discs (compare Fig. 7 A’ with Fig. 7 C’). Furthermore, in contrast to control discs, when we compared Ptc expression levels between glial cells adjacent to the MF and those in the posterior regions in mutant discs, we found that they were expressed at similar levels (compare purple arrows with red arrows in Fig. 7 D’). Thus, compared to control discs, the A/P ratio of Ptc levels was lower in discs with impaired cytoneme formation than in control glial cells (Fig. 7 G). Similar findings were observed when analyzing Ci expression levels (Fig. 7 E-F’’’ and H). Altogether, these results suggest that glial cytonemes are involved in Hh signalling transmission during eye disc development.

**Figure 7.**
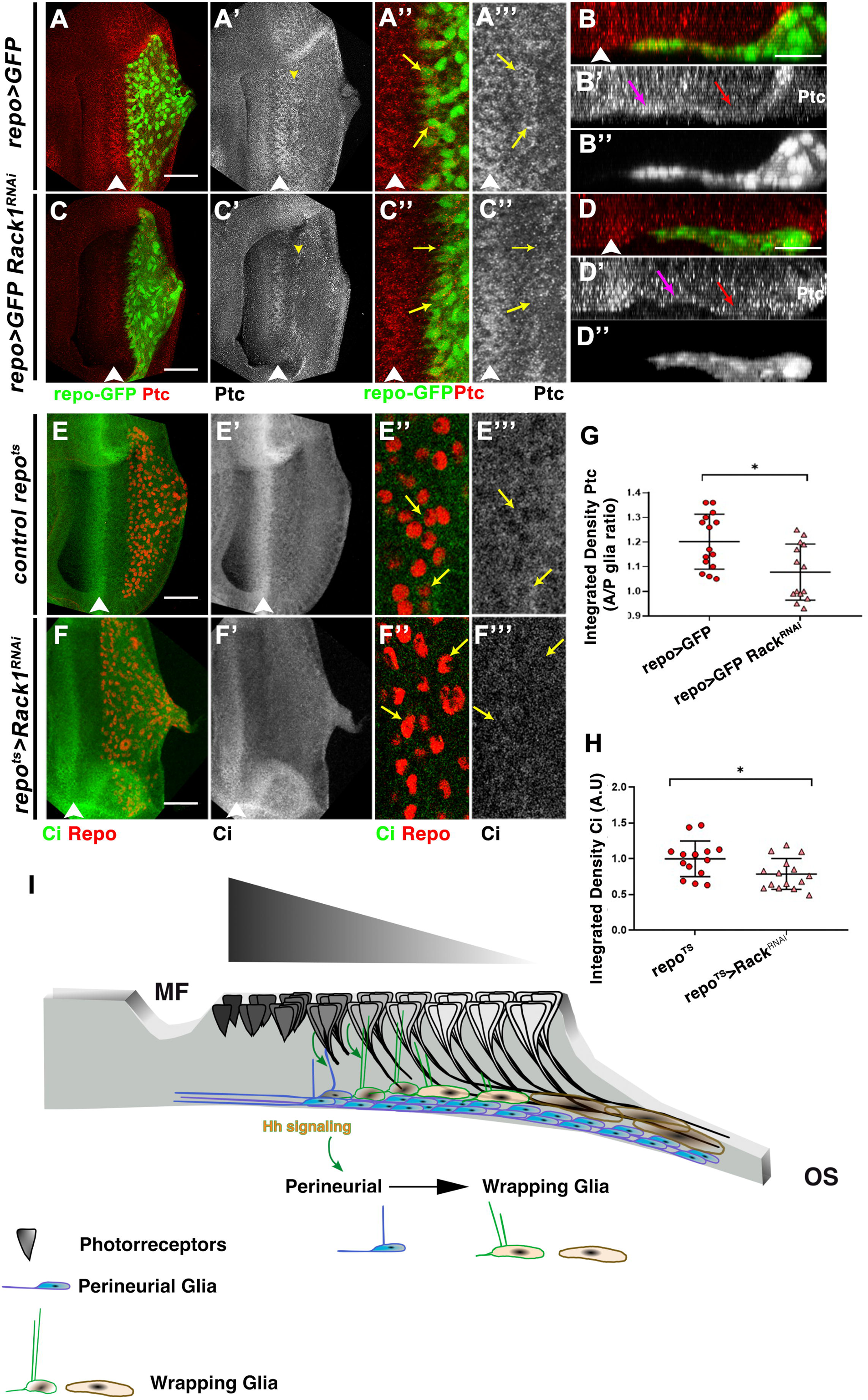
Inhibition of glial cytonemes results in a reduction of Hh signalling activation in glial cells. (A-D’’) *repo>GFP* (A-B’’) and *repo>GFP Rack1^RNAi^* (C-D’’) eye discs showing GFP (green) and Ptc (red). The MF is on the left (white arrowheads). Panels A’’, A’’’ and C’’, C’’’ are magnifications of A, A’ and C, C’, respectively. Panels B-B’’ and D-D’’’ are orthogonal views of the disc shown in A and B respectively. Ptc expression is higher in anterior glial cells (yellow arrows in A’’, A’’’, and magenta arrow in B’) compared to posterior glial cells (red arrow in B’). This difference is diminished with Rack1 depletion, with Ptc expression levels equalized across anterior and posterior glial cells (C’-D’’, magenta arrow in D’ vs. red arrow in D’). (E-F’’’) *repo>GFP* (E-E’’) and *repo>GFP Rack1^RNAi^* (F-F’’) eye discs stained for Ci (green) and Repo (red). Panels E’’, E’’’ and F’’, F’’’ are magnifications of E, E’ and F, F’, respectively. Yellow arrows indicate Repo-positive nuclei. Scale bars: 50 µm. (G) A/P ratio of Ptc integrated density and (H) normalized Ci integrated density of the described genotypes. *p<0.05. (I) Model: Glial cytonemes directed toward the MF are essential for receiving signals from MF cells. Additionally, glial cells extend cytonemes towards photoreceptors (PRs), activating Hh signaling. Hh is produced in a gradient by photoreceptors, with higher levels near the MF and lower levels posteriorly. PR-oriented cytonemes mediate Hh signaling activation, prompting precursor WG cells near the gradient to differentiate into mature WG cells.

We then investigated the effect of downregulating the Hh and Dpp signalling pathways in glial cells. To this end, we overexpressed Ptc, a negative regulator of the Hh pathway (reviewed in ^5^), and Brinker (brk), a transcriptional repressor of Dpp target genes^46^, both under the control of *repo-Gal4*. We observed that inhibition of Hh pathway in *repo-Gal4>UAS-ptc* discs only led to a mild reduction in glial cell number (Fig. S7 A-B’’’, D) and no effect on glial migration (Fig. S7 E). By contrast, inhibition of Dpp pathway in *repo-Gal4>UAS-brk* discs produces a much more pronounced reduction in glial cell number and migration (Fig. S7 A-A’’’, C-E). These effects are similar to those produced when cytonemes formation is impaired (Fig. 3). These results also suggest that Dpp pathway is more prominently involved in regulating glial cell number and migration than Hh, as it has been previously proposed ^21,23^.

Our observations also suggested that cytonemes played a critical role in the differentiation of WG cells. Since we found that glial cytonemes are targeted to cells producing Hh, and that Hh signalling is downregulated in glial cells when cytonemes are impaired, we next investigated whether disruption of these pathways affects WG differentiation. To investigate this, we blocked both pathways in WG cells by overexpressing *ptc* or *brk* using the *Mz97-Gal4* line. We found that downregulation of Hh signalling significantly reduced the number of WG cells and their proportion relative to the total glial population, whereas inhibition of Dpp had no significant effect (Fig. S8). Furthermore, the overexpression of a mutant form of *Ihog* (*Ihog-FN1\*\*\**), which impairs Hh signalling^47^, also produces a mild but statistically significant reduction in the proportion of WG cells (Fig. S8 C-C’’, E). All these results support the idea that cytonemes mediate the function of Hh signalling during WG differentiation.

Finally, we examined whether inhibiting Hh signalling in glial cells affects cytoneme formation and elongation. To assess this, we analysed the maximum length and orientation index of MF-oriented cytonemes in discs blocking the Hh pathway with the following genotypes: *repo-Gal4>UAS-Ihog-RFP UAS-ptc* and *repo-Gal4>UAS-Ihog-RFP UAS-Ci^RNAi^*. No significant differences were observed in these parameters compared to *repo-Gal4>UAS-Ihog-RFP* control discs (Fig. S9), suggesting that Hh signaling does not play a role in glial cytoneme formation. This independence of cytoneme formation from Hh signalling is consistent with previous data in wing discs and abdominal histoblasts ^33^.

## DISCUSSION

The development of a complex nervous system depends on precise interactions between neuronal and glial cells. Cell communication and intercellular signalling regulate the formation, specification, and survival of these cells, ensuring proper cell numbers and identities for functional neuronal networks. Cytonemes have emerged as a precise mechanism that facilitates the exchange of signals between cells (reviewed in ^29–32^), providing higher specificity than diffusion-based mechanisms and regulating key signalling pathways.

Cytonemes were first identified in the *Drosophila* wing imaginal disc^28^, where they mediate interactions between receptor and ligand-producing cells, enabling precise signalling over long distances. They also facilitate communication between epithelial wing disc cells, the tracheal air sac primordium (ASP) and the myoblasts (AMPs) that form the flight muscles ^29,48,49^. Cytonemes are also critical for asymmetric signalling and niche-specific organisation of AMPs^50^. This evidence highlights the role of cytonemes in mediating interactions between different cell types. In addition, cytonemes function as specialised signalling structures in vertebrates such as amphibians, chicks, zebrafish and mice, supporting the paracrine transport of key signalling molecules including Notch, Spi/EGF, Bnl/FGF, Dpp/BMP, Wg/Wnt and Hh/Shh ^29, 32^.

Although cytonemes play a critical role in mediating cellular interactions in various tissues, their involvement in coordinating glial and neuronal development during nervous system formation remains unexplored. This gap is likely due to the challenges of studying cytonemes in complex systems with multiple cell types and projections. Studies using the *Drosophila* eye disc have revealed fundamental mechanisms linking neuronal and glial development, positioning *Drosophila* glial cells as models for understanding mammalian glial biology^3–7,12,16^. The spatial separation of glial and neuronal populations in the eye disc provides a unique system for studying the distribution and function of cytonemes in coordinating glial and neuronal development.

### Glial migration is mediated by cytonemes, but they do not provide directional cues

During eye development, glial cells migrate from the optic stalk into the eye disc only after photoreceptor differentiation, independent of axonal substrates^18, 51^. Differentiating photoreceptors appear to regulate subretinal glial migration over distance, with Dpp and Hh signalling promoting glial motility^21,23^. Our results show that disruption of cytonemes alters glial migration and impairs Hh signalling, as evidenced by reduced expression of Hh target genes. While Hh acts as a strong motogen of glial movement, it does not provide directional cues or influence proliferation^21^. Consistently, disruption of these ses does not affect glial proliferation or their preference to migrate towards the morphogenetic furrow (Fig. 3 and Fig. S4).

These findings suggest that glial cell cytonemes mediate the activation of the Hedgehog (Hh) signalling pathway in response to photoreceptor-derived Hh. Disruption of cytonemes impairs Hh signalling and alters glial migration. Motility analysis revealed that cytonemes disruption reduced angular entropy compared to controls, where higher entropy typically indicates random or uncoordinated movement. Although counterintuitive, this may reflect the complexity of intercellular signalling. In control conditions, competing signals from multiple photoreceptors could increase angular entropy by pulling cells in different directions. Conversely, cytonemes disruption may limit intercellular interactions while preserving positional cues, leading to more directional movement and reduced entropy.

Disrupting cytonemes impairs glial migration, underscoring their importance in facilitating interactions necessary for coordinated glial and neuronal development. While Hh influences glial migratory behaviour without affecting proliferation, Dpp promotes both proliferation and motility of subretinal glia ^21,23^. The effects of cytonemes disruption closely resemble those of impaired Hh signaling, which primarily affects motility. However, given cytonemes established role in mediating Dpp signaling in various tissues, it is plausible that cytonemes disruption also impacts Dpp signaling. Notably, inhibiting Dpp signaling causes a more pronounced defect in glial migration than Hh signaling (Fig. S7). Residual Dpp activity may remain functional after cytoneme disruption, sufficient to support proliferation but inadequate to fully sustain motility. Thus, cytonemes disruption likely affects both Hh and Dpp pathways, altering glial motility.

### Wrapping glia differentiation

For a functional eye, glial subtypes must match the number of photoreceptor neurons. During development, the differentiation of perineurial glia into wrapping glia cells is tightly regulated to maintain the correct ratio of wrapping glia to photoreceptor axons. FGF receptor signalling is critical for glial development in the eye disc, as its downregulation reduces glial cell numbers, disrupts migration and impairs differentiation^3^.

Drosophila has two FGF receptor genes^52,53^, one of which, *heartless* (*htl*), is expressed in embryonic and eye disc glial cells, with highest levels at the leading edge of migrating glia^3,54^. Activation of Htl depends on two FGF-8-like ligands, Pyramus and Thisbe, which regulate its function^3,55–57^. Pyramus probably acts as an autocrine signal that stimulates glial cell division. In addition, its expression in the stripe of cells anterior to the morphogenetic furrow suggests a role in carpet cell outgrowth and glial migration towards the morphogenetic furrow^58^. Contact between wrapping glial precursors and nascent photoreceptors expressing Thisbe induces a peak in Htl activity, triggering differentiation of perineurial glia into wrapping glia^59^. High FGF signalling activates the transcription factor Cut, which is essential for wrapping glia differentiation^59^, whereas low FGF signalling supports proliferation and migration^4,6^.

In this work we have identified photoreceptor-oriented cytonemes produced by glial cells that contact nascent photoreceptors. While these cytonemes are clearly observed in wrapping glia, we also found that some perineurial glial cells can produce them. It is possible that perineurial progenitor cells produce these cytonemes in an attempt to contact nascent photoreceptors. Once in contact, the cytonemes might stabilise, and this stabilisation could be crucial for promoting the activation of the Hh signaling pathway in glial cells, facilitating their differentiation into the WG (Fig. 7I).

The photoreceptor-oriented cytonemes make contact with photoreceptors that produce the Hh ligand, and we have shown that this pathway is essential for the differentiation of WG. Given the critical role of FGF signalling in glial development and differentiation, it is plausible that both the Hh and FGF signalling pathways interact synergistically in this process. Furthermore, it is possible that these photoreceptor-oriented cytonemes also play a role in mediating FGF signalling during WG differentiation. We propose several potential scenarios to further investigate this interaction.

One possibility is that certain photoreceptor-oriented cytonemes in glial cells contain the Htl receptor. Interaction with photoreceptors expressing the ligand Thisbe enhances FGF signaling, driving glial differentiation into WG. During tracheal air sac primordium (ASP) development, ASP cells extend cytonemes containing specific receptors like Btl (FGF receptor) to contact Bnl (FGF)-expressing cells or Tkv (Dpp receptor) to interact with Dpp-expressing cells, but not both simultaneously^48^. This highlights the existence of signal-specific cytonemes mediating FGF signaling. Similarly, cooperation between FGF-activating and Hh-mediated cytonemes may be crucial for WG differentiation.

It has been observed that cytonemes are present in adult muscle precursors (AMPs) within wing discs, and that they mediate FGF signalling. AMPs proliferate to generate transient amplifying cells that undergo myogenic fusion and differentiation into adult flight muscles^60,61^. Polarised cytonemes extended by AMPs contact epithelial junctions in FGF-producing regions via FGFR/Heartless (Htl), allowing direct FGF signal reception^50^. The *htl* gene knockdown in AMPs resulted in the disruption of cytonemes targeting the disc, while lateral cytonemes remained unaffected. This suggest that FGF signalling may be essential for maintaining the stability of cytonemes^50^. It is therefore possible that a cooperative mechanism between Hh and FGF signalling may involve FGF activation to stabilise photoreceptor-oriented cytonemes, which would then facilitate effective Hh signalling and promote WG differentiation.

Finally, in the ganglionic branches of the tracheal system, Hh signalling has been shown to upregulate Bnl/FGF levels^62^. This implies that PR-oriented cytonemes activating Hh signalling could enhance FGF signalling, ensuring proper WG differentiation.

In summary, our results suggest a model in which glial cells produce two distinct sets of cytonemes, each based on their spatial orientation. The first set, oriented towards the MF, likely participates in receiving signals from the MF, such as Dpp. The second set, directed towards nascent photoreceptors, is involved in receiving Hh signal and promoting the differentiation of certain perineurial glial cells into WG (Fig. 7 I). Through the regulation of these signalling pathways, glial cytonemes control both the migration and differentiation of glial cells—two essential aspects of glial behaviour that are closely linked to neuronal differentiation. Thus, cytoneme-mediated signalling appears to play a critical role during the coordinated development of glial cells and neurons in the *Drosophila* eye disc. Since this mechanism may be conserved across species, our findings could also offer insights into signalling regulation during vertebrate nervous system development.

## RESOURCE AVAILABILITY

### Lead Contact

The data and materials supporting the findings of this study are available upon request to. Interested parties may contact the corresponding author to obtain access to the data and materials for further examination and verification.

## ACKNOWLEDGMENTS

We thank Carlos Estella for providing reagents and useful discussion. We would like to thank Ana Citlali and Adrián Aguirre for their invaluable help with cytonemes quantification and for generously providing fly stocks. We are also very grateful to Ginés Morata for his constant support throughout the development of this project. We are very grateful to the Bloomington Stock Center and the Developmental Studies Hybridoma Bank for providing fly strains and antibodies. This study was supported by grants from: FEDER/ Ministerio Ciencia-Agencia Estatal de Investigación PID2020-114533GB-C21, PID2021-127114NB-I00 and institutional grant from Banco de Santander to the CBMSO. None of the authors have any competing interest.

## AUTHOR CONTRIBUTIONS

J.M.G-A performing experiments, analysis data, designing experiment, writing and editing text; G.G performing experiments, analysis data, Generating the code; D.F. performing experiments, analysis data, writing and editing text; I.G. planning experiment, writing and editing text and A.B., designed the experiments, discussed, analyzed data, and prepared the manuscript

## METHODS

### *Drosophila* stocks and genetics

All *Drosophila* strains used on this study have been maintained with standard medium and at 25°C. Crosses were maintained at 17°C before the inhibition of Gal80^ts^ at 29°C.

The following stocks were used:

Gal4 lines: *repo-Gal4* (BDSC# 7415), *tub-Gal80^ts^*^63^, *Mz97-Gal4, C527-Gal4*^20^*, GMR-Gal4*^64^.

*UAS lines: UAS-Ihog-RFP*^65^, *UAS-Moe-Cherry*^66^, *UAS-Ihog-CFP*^34^*, UAS-Lifeact-mRFPruby* (BDSC#58362)^35^, *UAS-Rack1^RNAi^* (BDSC#34694), *UAS-cora^RNAi^* (VDRC#v9787), *UAS-pico^RNAi^* (VDRC#v16371), *UAS-GFP* (II and III) (described in Flybase, Bloomington Drosophila Stock Centre), *UAS-ptc*^67^, *UAS-brk*, *UAS-IhogΔFN1-RFP* is a mutant form of Ihog, containing three mutual mutations^47^, *UAS-Ci^RNAi^* (NIG-FLY, 2125R-1).

Reporter lines: *BAC-Hh:GFP*^41^, *dpp-LacZ* (p10638, BDSC#12379). Other lines: *repo-RFP*^39^.

### Imaginal discs staining

Third instar larvae were dissected in PBS1X on ice and fixed with 4% paraformaldehyde, 0.1% deoxycholate (DOC) and 0.1% Triton X-100 for 27 min at RT. Then they were washed with a blocking solution (PBS1X, 1% BSA, and 0.3% Triton X-100) and incubated with the corresponding primary antibody over night at 4 °C. After that, they were washed in PBS1X, 0.3% Triton X-100 and incubated with the corresponding fluorescent secondary antibodies for at least 2 hours at RT in the dark. They were washed and mounted in Vectashield mounting medium (Vector Laboratories).

The following primary antibodies were used: mouse anti-Repo (Developmental Studies Hybridoma Bank, DSHB#8D12) 1:25^68^; rat anti-Elav (DSHB# 7E8A10) 1:25; rabbit anti-Dcp1 (Cell Signal Technology) 1:200; rabbit anti-PH3 (Cell Signal Technology) 1:200; rat anti-Ci (DSHB#2A1) 1:50^69^; mouse anti-b-galactosidase (DSHB#40-1a) 1:50; mouse anti-ptc (DSHB#Apa1) 1:50^70^. Fluorescently labelled secondary antibodies (Molecular Probes Alexa-488, Alexa-555, Alexa-647, ThermoFisher Scientific) 1:200.

### *In vivo* imaging of eye discs

Third instar larvae from the appropriate genotype were washed in 1× PBS and dissected in Schneider’s medium (11720-034, Invitrogen), supplemented with 15% fetal bovine serum (10106-169, Invitrogen) and 0.5% penicillin-streptomycin (15140-122, Invitrogen). Eye discs were carefully separated from the CNS with entomological needles and transferred to 35 mm glass-bottom culture dishes coated with poly-D-lysine (P35GC-1.5-10-C, Mattek Corporation) with more supplemented Schneider’s medium and methylcellulose (M0387-100G, Sigma-Aldrich) to a final concentration of 0.3% to maintain the discs in place. Time-lapse movies were obtained using an inverted confocal microscope (IX83 Olympus) coupled to a confocal spinning disk SpinSR10 (Olympus) at room temperature or 29°C. Stack images were taken every 2 min for 2-6h

### Temperature shift experiments

Larvae of genotypes *repo^ts^-Gal4>UAS-Ihog-RFP UAS-Rack1^RNAi^, repo^ts^-Gal4>UAS-Ihog-RFP UAS-cora^RNAi^*, *repo^ts^-Gal4> UAS-Ihog-RFP UAS-Pico^RNAi^* and *GMR^ts^>UAS-Ihog-RFP UAS-Rack1^RNAi^* were maintained at 17°C until L3 stage and raised at a restrictive temperature of 29°C for 24 hours before dissecting them to allow for the degradation of the Gal80^ts^ factor.

### Image acquisition, quantifications and statistical analysis

Stack images were captured with a Nikon A1R and a Leica (Solms, Germany) DB550 B and a Spinning Disk SpinSR10 confocal microscopes. Multiple focal planes were obtained for each imaginal disc. Quantifications and image processing were performed using the Fiji/ImageJ (https://fji.sc) and Adobe Photoshop software.

Glial index was calculated as the ratio between the total number of glial cells, detected by positive Repo staining, and the area of the region occupied by photoreceptor cells, detected by positive Elav staining. In a similar manner, WG index was calculated as the ratio between the total number of WG cells, detected by positive GFP signal, and the size of the region occupied by photoreceptor cells. For the WG proportion (%WG), the number of WG cells was divided between the number of total glial cells. Glial migration index was measured as the ratio between the number of photoreceptor rows behind the MF that reach the most anterior glial cells and the total number of rows in the disc.

Maximum MF-oriented cytonemes length was calculated as the mean of the five longest cytonemes per disc, by using the “Straight Line” tool in ImageJ, from the tip to the end of the cytoneme. The same procedure was used to measure the maximum length of PR-oriented cytonemes, retinal and WG cytonemes. MF-cytoneme index was calculated as the ratio between the total number of MF-oriented cytonemes, using the “Multi Point” tool in ImageJ, and the length of de disc occupied by Ihog-RFP expression, measured by the “Straight Line” tool in ImageJ. To calculate the PR-cytoneme index, an orthogonal section was first made, using the “Reslice” tool in ImageJ, in the area corresponding to the most anterior glia. Then, the procedure was performed in the same way than described above for the MF-cytoneme index. All orthogonal sections were made with an output spacing of 0,5 µm and 40 slices.

Apoptotic index was measured as the ratio between the Dcp1 positive area in the region of the disc occupied by glial cells, selected by the “Treshold” tool in ImageJ, and the total number of glial cells. We measured mitotic index as the ratio between the number of PH3-positive particles in glial cells and the total number of glial cells (particle size was >5µm).

For calculating the A/P ratio Ptc, the mean intensity of Ptc signal was measured in the most anterior glial cells and divided between the mean intensity of Ptc signal in the most posterior glial cells, selected by a region of the same size. Ci mean intensity was also calculated in the area occupied by glial cells, and normalized respect to *repo-Gal4* control discs.

Statistical analyses were performed using the GraphPad Prism software (https://www.graphpad.com). To compare between two groups, a non-parametric Student’s *t*-test test was used. To compare between more than two groups, a non-parametric, one-way ANOVA Dunnett’s test was used (in Fig. S6 and S7). Sample size was indicated in each Figure.

### Tracking Data Collection

#### Preprocessing

In order to compensate for eye disc drift during *in vivo* capture, we stabilized the video to gather accurate data. Sample drift was corrected using plugin Hyperstackreg, which compensates for both rotation and translation by applying a transformation matrix. Additionally, depending on the sample, an affine transformation was performed to further adjust the drift.

#### Data Gathering

We tracked glial cell movements using TrackMate2^70,71^. Cells were automatically identified using the StarDist (3) segmentation algorithm, and tracks were generated using the LAP tracker^72^, adjusting the frame skip based on sample-specific conditions.

#### Data filtering

Low-quality tracks were filtered based on their median prediction fidelity or if the tracking prediction between frames fell below a sample-adjusted threshold.

### Data Analysis

#### Entropy

For the movement analysis, we calculated the movement vector between successive frames using the following formula.

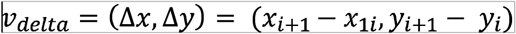

The angle, representing the orientation of each movement vector relative to the x-axis, was computed using the formula.

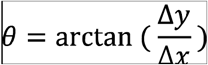

After that, the angular difference between two consecutive movement vectors was calculated using the formula

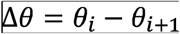

The interval of all possible angle values 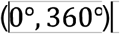 was divided into 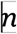 bins, and the number of angles between consecutive vectors 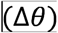 in each bin was counted to create a frequency diagram. The optimal number of bins for entropy calculations was determined using the Friedman-Diaconis rule:

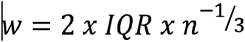

Following a simplified version of ^38^ we quantified the variability of the frequency distribution using the Shannon entropy formula to assess the complexity of the movement patterns:

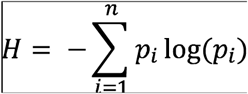

Furthermore, since 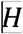 is correlated with the number of bins 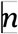, the values were normalized by dividing by the maximum possible entropy for a given number of bins. This ensured that entropy values ranged from 0 (indicating more ballistic movement) to 1 (indicating more chaotic movement).

### Cytonemes analysis

#### Cytonemes lifespan

Based on ^73^ we calculated the lifespan per cytoneme as the number of frames between the appearance of a cytoneme and the retraction of the same cytoneme and divided it by the unit of time per frame:

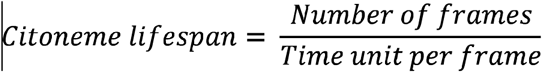

#### Cytonemes Parallelism

After gathering the base 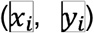 and end 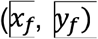 coordinates of each cytoneme, a vector was created for each. The parallelism was then determined by calculating the angle between all vectors (cytonemes) within each sample using the following formulas.

#### 1. Angle Calculation

For each cytoneme, the angle 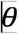 between the cytoneme vector and the x-axis is calculated using:

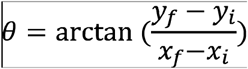

Where:

- 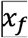 and 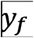 are the coordinates of the cytoneme’s end point.
- 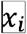 and 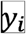 are the coordinates of the cytoneme’s base point.

#### 2. Pairwise Angle Differences

After calculating the angles for each cytoneme, the pairwise absolute differences in angles between each pair of cytonemes are computed as:

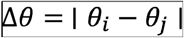

Where 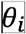 and 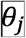 are the angles of two different cytonemes, and the result is converted from radians to degrees by multiplying by 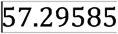 (since 1 radian ≈ 57.2958 degrees).

