## Supplementary Legends for "Cytoneme-mediated signalling coordinates the development of glial cells and neurons in the *Drosophila* eye"

**Figure S1. *Ihog-RFP* expression in glial or retinal cells does not affect glial cell number or migration. (**A) Glial index and (B) glial migration of control (wt), *repo>Ihog-RFP* and *GMR>Ihog-RFP* eye discs showing no statistical differences in glial number or migration. ns, not significant. C) Graph comparing between MF-oriented and PR-oriented maximum cytoneme length in *repo>Ihog-RFP* discs (n=21 and 19, respectively).

**Figure S2. Glial and retinal cytonemes are actin-based structures. (**A, A') Magnification of the basal region of a *repo>Moe-Cherry* eye disc showing Repo (green), *Moe-Cherry* (red) and Elav (blue) staining. Glial cells form cytonemes (yellow arrow) towards the morphogenetic furrow (MF, white arrowhead). (B, B') Orthogonal view of the disc shown in A (white dashed line in A), optic stalk to the right. Glial cytonemes are oriented towards the MF (yellow arrow) and photoreceptor cells (purple arrow). (C-D') Orthogonal views in the anterior (C, C', yellow dashed line in A) and in the posterior (D, D', blue dashed line in A) regions of the disc shown in A. Photoreceptor oriented cytonemes are only produced by the most anterior glia (compare C, C' with D, D'). (E, E') High magnification of disc shown in A, A' revealing MF-oriented cytonemes (yellow arrows). (F, F') High magnification of C, C' showing photoreceptor oriented cytonemes. (G, G') Magnification of an *GMR> Moe-Cherry* eye disc showing Repo (green), *Moe-Cherry* (red) and Elav (blue) staining. Retinal cells produce cytonemes (yellow arrow) oriented towards the MF (white arrowhead). (H, H') Orthogonal view of the disc shown in G (white dashed line in G), optic stalk to the right. Retinal cells produce MF oriented cytonemes (yellow arrow) in the basal region of the disc. Orthogonal views in the anterior (I, I', yellow dashed line in G) and posterior (J, J', blue dashed line in G) regions of the disc shown in G. No glial oriented cytonemes are visible. (K, K') High magnification of G, G'. (L, L') High magnification of H, H'. In both cases MF-oriented retinal cytonemes are shown. Scale bars, 20μm.

**Figure S3. Inhibition of glial cytonemes reduces their number and maximum length.** Magnifications of *repo$^{ts}$>Ihog-RFP* (A, A'''), *repo$^{ts}$>Ihog-RFP Rack1$^{RNAi}$* (B, B'''), *repo$^{ts}$>Ihog-RFP cora$^{RNAi}$* (C, C''') and *repo$^{ts}$>Ihog-RFP pico$^{RNAi}$* (D, D''') eye discs staining for Repo (green), Ihog-RFP (red) and Elav (blue). MF is on the left (white arrowhead). Panels in A”-A’”, B”-B’”, C”- C’”, and D”-D’” correspond to orthogonal views of the anterior regions of discs in A, B, C, and D, respectively. Scale bars, 20μm. MF-oriented cytoneme length (E), MF-oriented cytoneme index (F) and photoreceptor-oriented cytoneme index (G) of discs of the genotypes described above. **p<0,01, ***p<0,001, ****p<0,0001.

**Figure S4. Reduction of glial cell number after disrupting glial cytoneme is not due to apoptosis or proliferation.** *repo$^{ts}$>Ihog-RFP* (A, A’), *repo$^{ts}$>Ihog-RFP Rack1$^{RNAi}$* (B, B’) and *repo$^{ts}$>Ihog-RFP cora$^{RNAi}$* (C, C’) eye discs showing Dcp1 (green) and Repo (red) staining. Glial areas are marked with a yellow line. (D) Apoptotic index of the genotypes described above. *repo$^{ts}$>Ihog-RFP* (E, E’), *repo$^{ts}$>Ihog-RFP Rack1$^{RNAi}$* (F, F’) and *repo$^{ts}$>Ihog-RFP cora$^{RNAi}$* (G, G’) eye discs staining for PH3 (green) and Repo (red). Glial areas are marked with a yellow line. (H) Mitotic index of the genotypes described above. ns, not significant. Scale bars, 50μm.

**Figure S5. Retinal cytonemes inhibition reduces their number and maximum length.** Magnifications of *GMR$^{ts}$>Ihog-RFP* (A-A’), *GMR$^{ts}$>Ihog-RFP Rack1$^{RNAi}$* (B-B’’) eye discs showing Ihog-RFP (green) and Elav (blue) staining. Scale bars, 20μm. MF-oriented cytoneme length (C) and MF-cytoneme index (D) of discs of the genotypes described above. ****p<0,0001.

**Figure S6. Inhibition of retinal cytonemes has no major effect on glial cell number and migration.** (A-B’’’) Eye discs of genotypes *GMR$^{ts}$>Ihog-RFP* (A-A'''), *GMR$^{ts}$>Ihog-RFP Rack1$^{RNAi}$* (B-B''') showing Repo (green) and Elav (blue) staining. Panels in A'', A''', B'' and B''' correspond to magnifications of A, A' and B, B', respectively. MF is on the left (white arrowhead). The white dashed lines in A'' and B'' represent the distance between the most anterior glial cell and the first row of photoreceptors. Scale bars, 50 μm. (C) Glial index and (D) glial migration index of discs from the genotypes described above. ns, not significant; *p<0.05.

**Figure S7. Inhibition of Hh and Dpp signalling in glial cells reduces glial cell number and impairs glial migration in the eye disc.** *repo-Gal4* (A-A'''), *repo>ptc* (B-B''') and *repo>brk* (C-C''') eye discs showing Repo (green) and Elav (blue) staining. MF is to the left (white arrowheads). Panels in A'', A''', B'', B''', C'', C''' represent magnifications of A, A', B, B', C and C', respectively. The white dashed lines in A'', B'' and C'' represent the distance between the most anterior glial cell and the first photoreceptor row. Scale bars, 50μm. Glial index (D) and migration index (E) of discs from the genotypes described above. ns, not significant; *p<0,05; ****p<0,0001.

**Figure S8. Inhibition of Hh signalling in wrapper glial cells leads to impaired glial differentiation**. *Mz97>GFP* (A-A'''), *Mz97>GFP ptc* (B-B'''), *Mz97>GFP IhogFN**** (C-C'') and *Mz97>GFP brk* (D-D''') Eye discs showing GFP (green), Repo (red) and Elav (blue) staining. MF is on the left (white arrowheads). Scale bars, 50μm. (E) %WG of discs from the above genotypes. ns, not significant; *p<0.05; ****p<0.0001.

**Figure S9. Hh signalling is not involved in glial cytoneme formation.**
Magnifications of *repo>Ihog-RFP* (A, A'''), *repo>Ihog-RFP ptc* (B, B'''), and *repo>Ihog-RFP ci$^{RNAi}$* (C, C''') eye discs showing Repo (green), and Ihog-RFP (red) staining. MF is on the left (white arrowhead). Scale bars, 20μm. MF-oriented cytoneme length (D) and MF-oriented cytoneme index (E) of discs from the genotypes described above. ns, not significant.

**Video 1. 3D reconstruction of a *repo>Ihog-RFP* third instar eye disc**
This video shows a 3D reconstruction of a *repo>Ihog-RFP* third instar eye disc, highlighting the presence of cytonemes. The staining shows Repo (green), Ihog-RFP (red), and Elav (blue). Glial cells are observed extending cytonemes both toward the morphogenetic furrow and nascent photoreceptors.

**Video 2. Ihog-labelled cytonemes dynamics.**

Time-lapse imaging of an eye disc of genotype *repo>Ihog-RFP* showing an amplification of MF-oriented glial cytonemes. Timeframes of 2 minutes are shown of a total duration of 2 hours. Related to Fig.2.

**Video 3. Life actin-labelled cytonemes dynamics.**
Time-lapse imaging of an eye disc of genotype *repo>Lifeact-mRFPruby* showing an amplification of MF-oriented glial cytonemes. Timeframes of 2 minutes are shown of a total duration of 2 hours. Related to Fig.2.

**Video 4. Glial motility in control discs.**
Time-lapse imaging of an eye disc of genotype *repo-RFP* showing glial cells nuclei. Timeframes of 2 minutes are shown of a total duration of 2 hours. Related to Fig.5.

**Video 5. Glial trajectories in control discs.**
Time-lapse imaging of an eye disc of genotype *repo-RFP* showing all glial trajectories within the disc tracked with TrackMate2. Timeframes of 2 minutes are shown of a total duration of 2 hours. Related to Fig.5.

**Video 6. Glial motility under cytoneme disruption.**
Time-lapse imaging of an eye disc of genotype *repo-RFP repo>Rack1*$^{RNAi}$ showing glial cells nuclei. Timeframes of 2 minutes are shown of a total duration of 2 hours. Related to Fig.5.

**Video 7. Glial trajectories under cytoneme disruption.**
Time-lapse imaging of an eye disc of genotype *repo-RFP repo>Rack1*$^{RNAi}$ showing all glial trajectories within the disc tracked with TrackMate2. Timeframes of 2 minutes are shown of a total duration of 2 hours. Related to Fig.5.

**Video 8. WG differentiation in control discs.**
Time-lapse imaging of an eye disc of genotype *repo-RFP Mz97>GFP* showing total glial cell population in red and WG in green. White arrows indicate glial cells progressively acquiring GFP and differentiating into WG. Timeframes of 2 minutes are shown of a total duration of 146 minutes. Related to Fig.8.

**Video 9. WG differentiation under cytoneme disruption.**

Time-lapse imaging of an eye disc of genotype *repo-RFP Mz97>GFP Rack1*$^{RNAi}$ showing total glial cell population in red and WG in green. White arrows indicate glial cells progressively losing GFP. Timeframes of 2 minutes are shown of a total duration of 146 minutes. Related to Fig.8.
