## Supplementary figures and images for "Cytoneme-mediated signalling coordinates the development of glial cells and neurons in the *Drosophila* eye"

### Fig s1

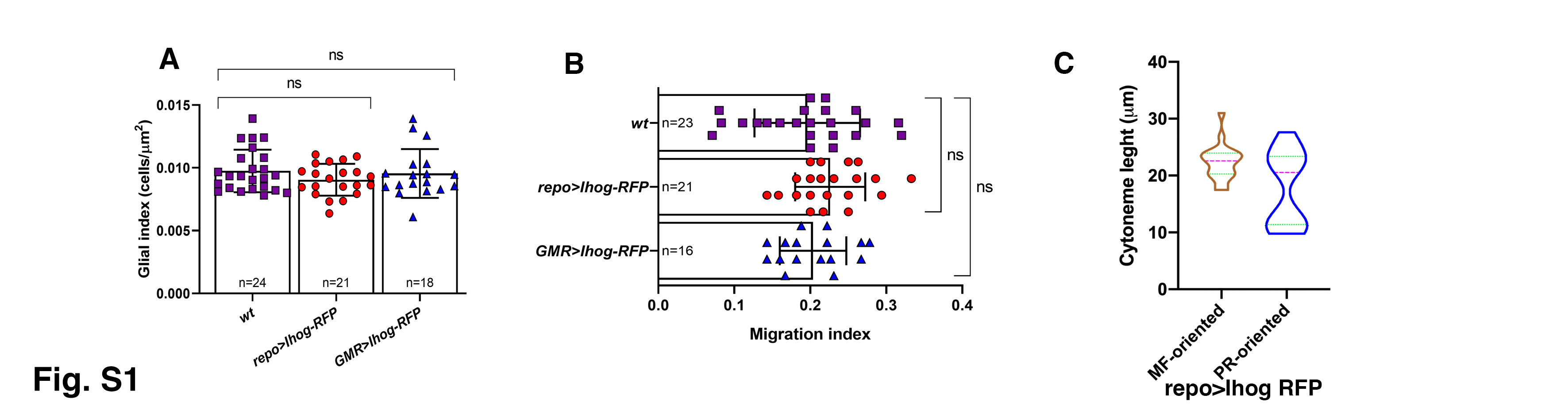

### Fig S2

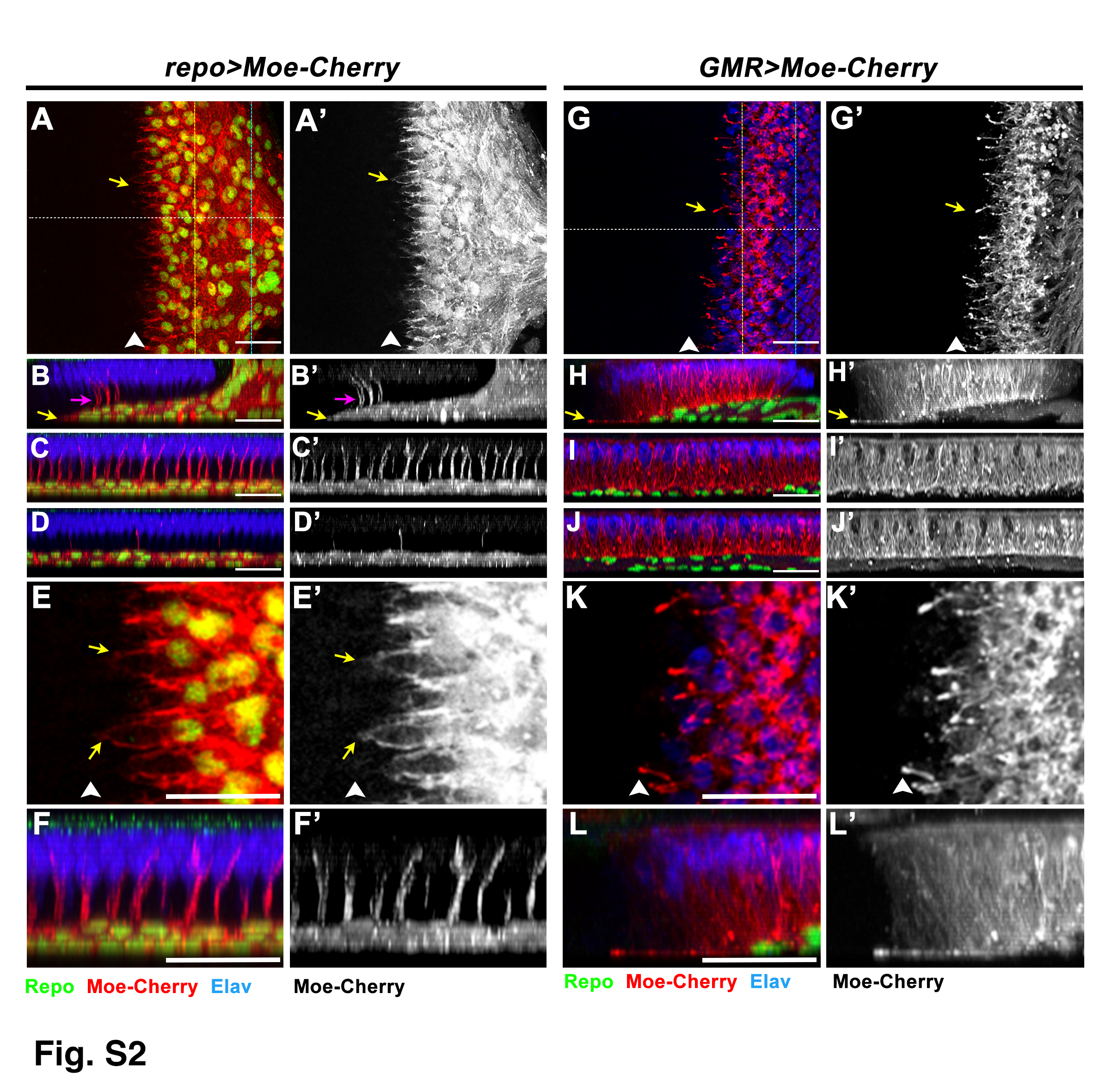

### Fig S3

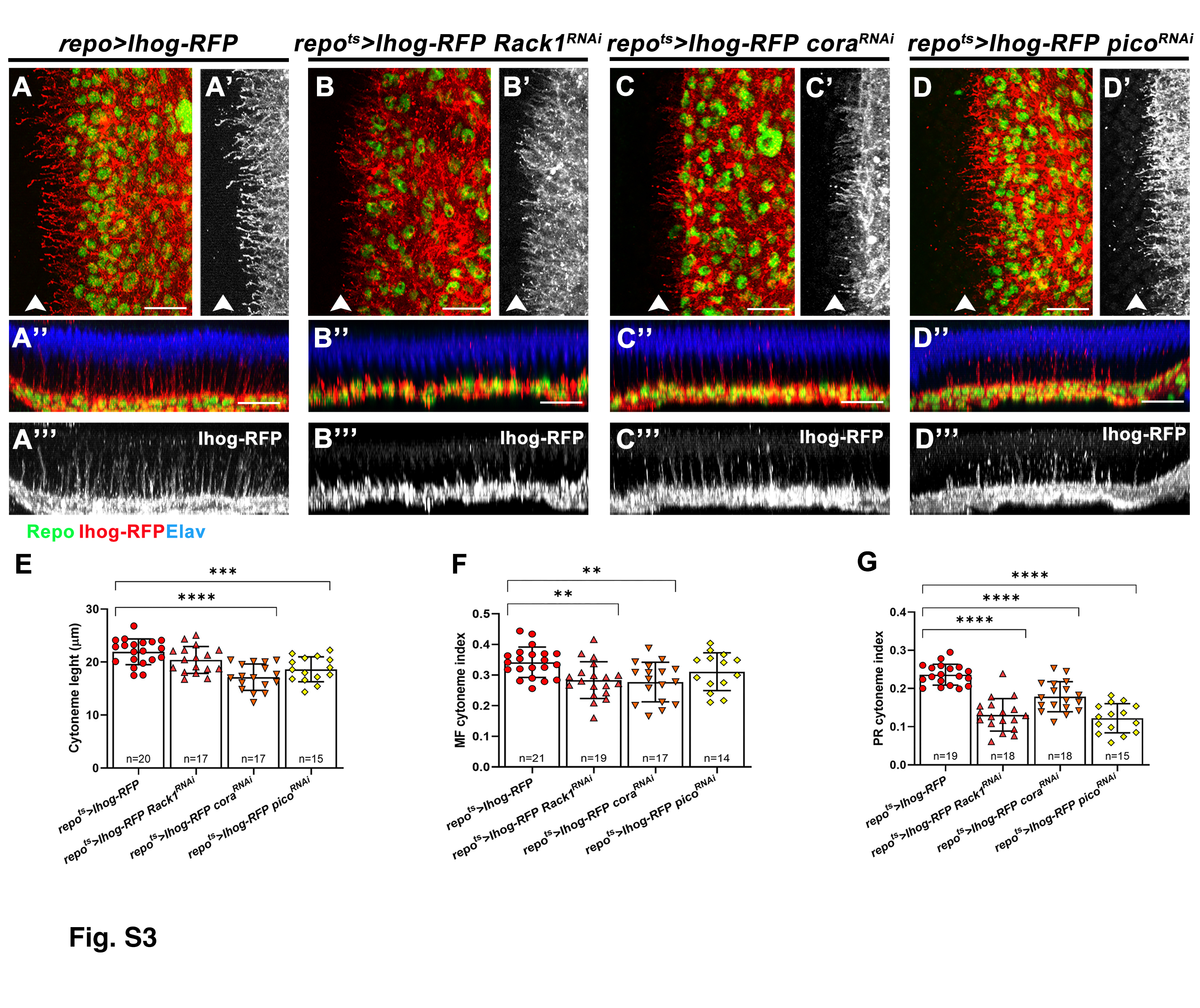

### Fig S4

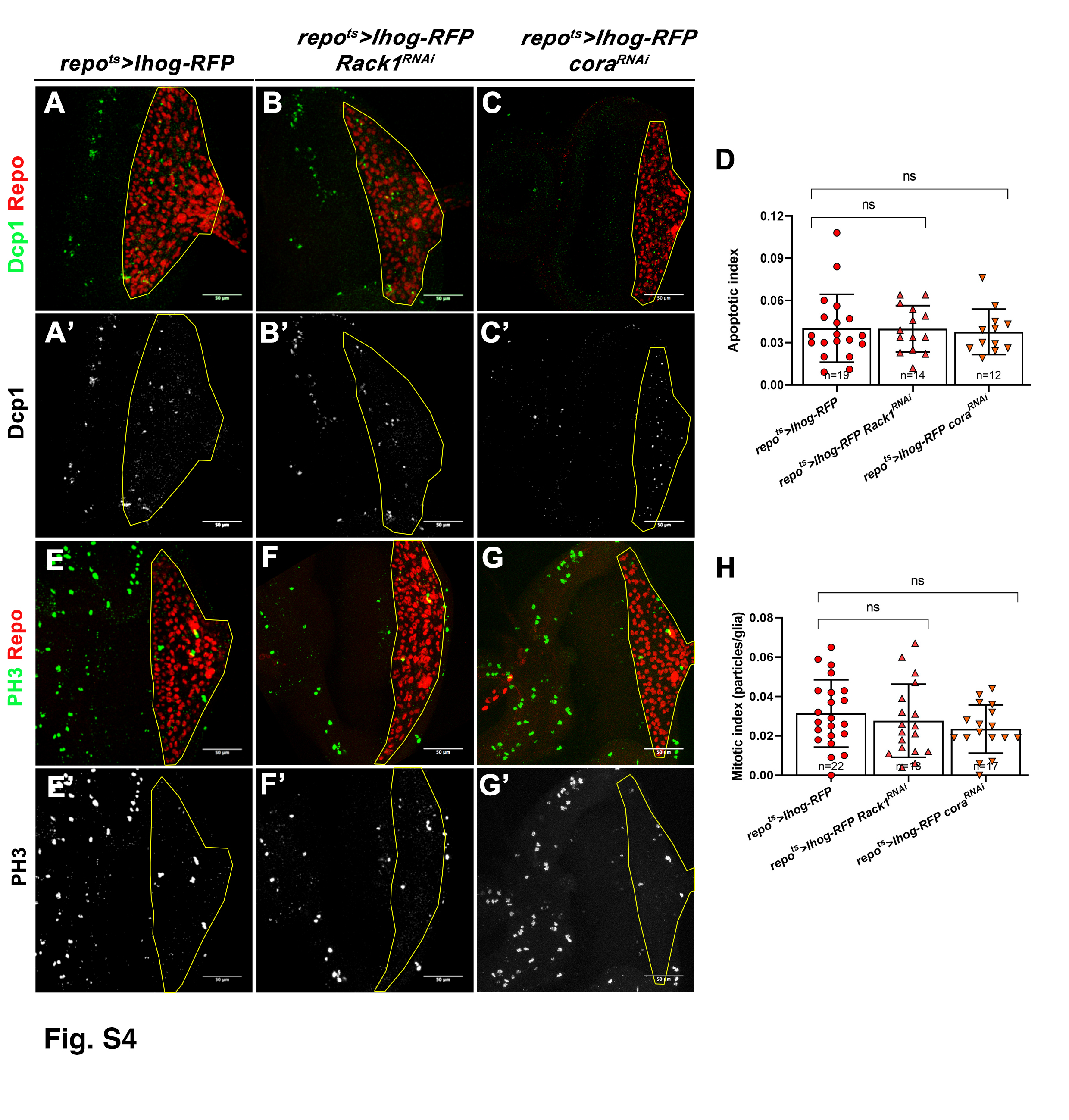

### Fig S5

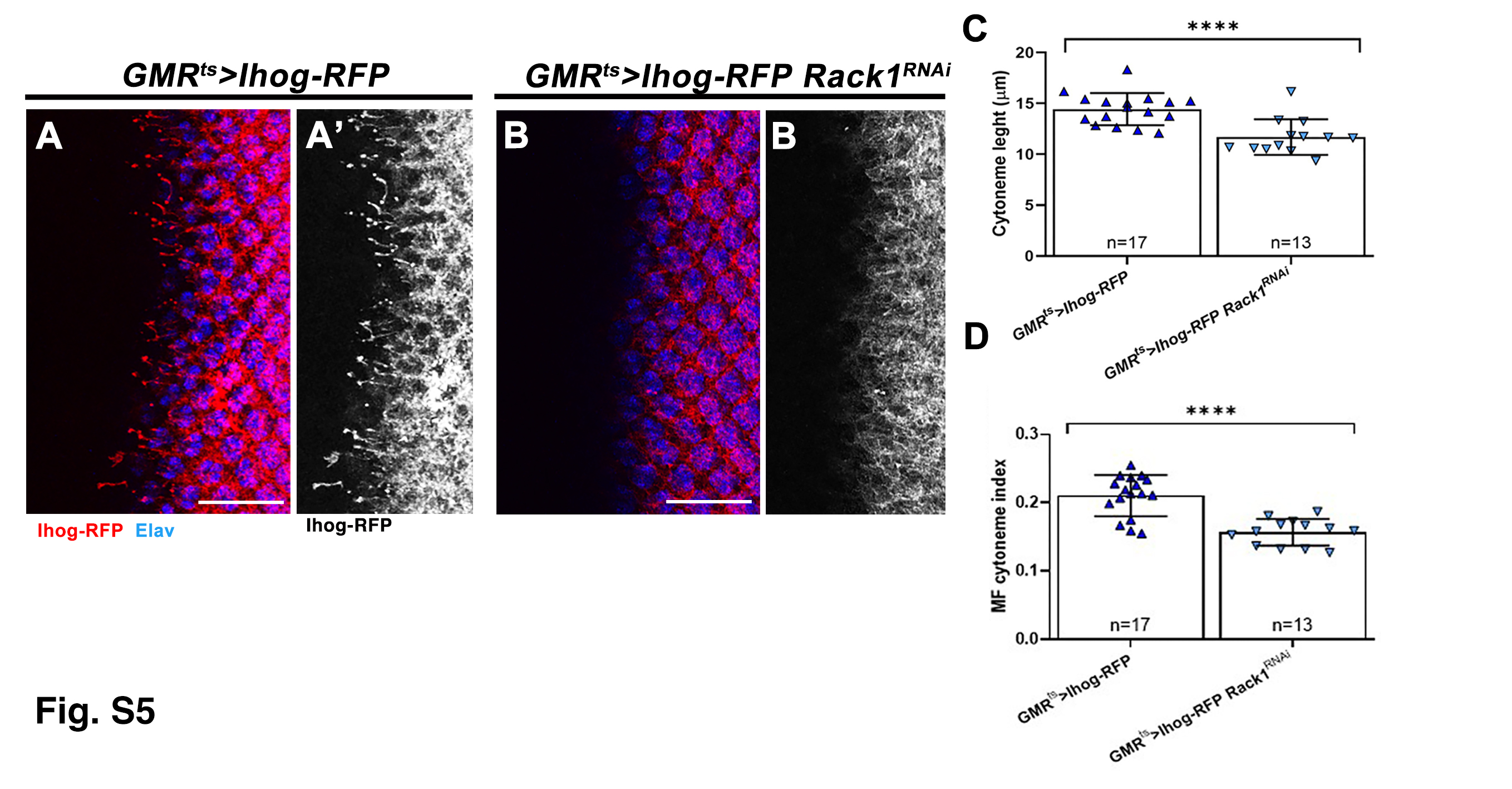

### Fig S6

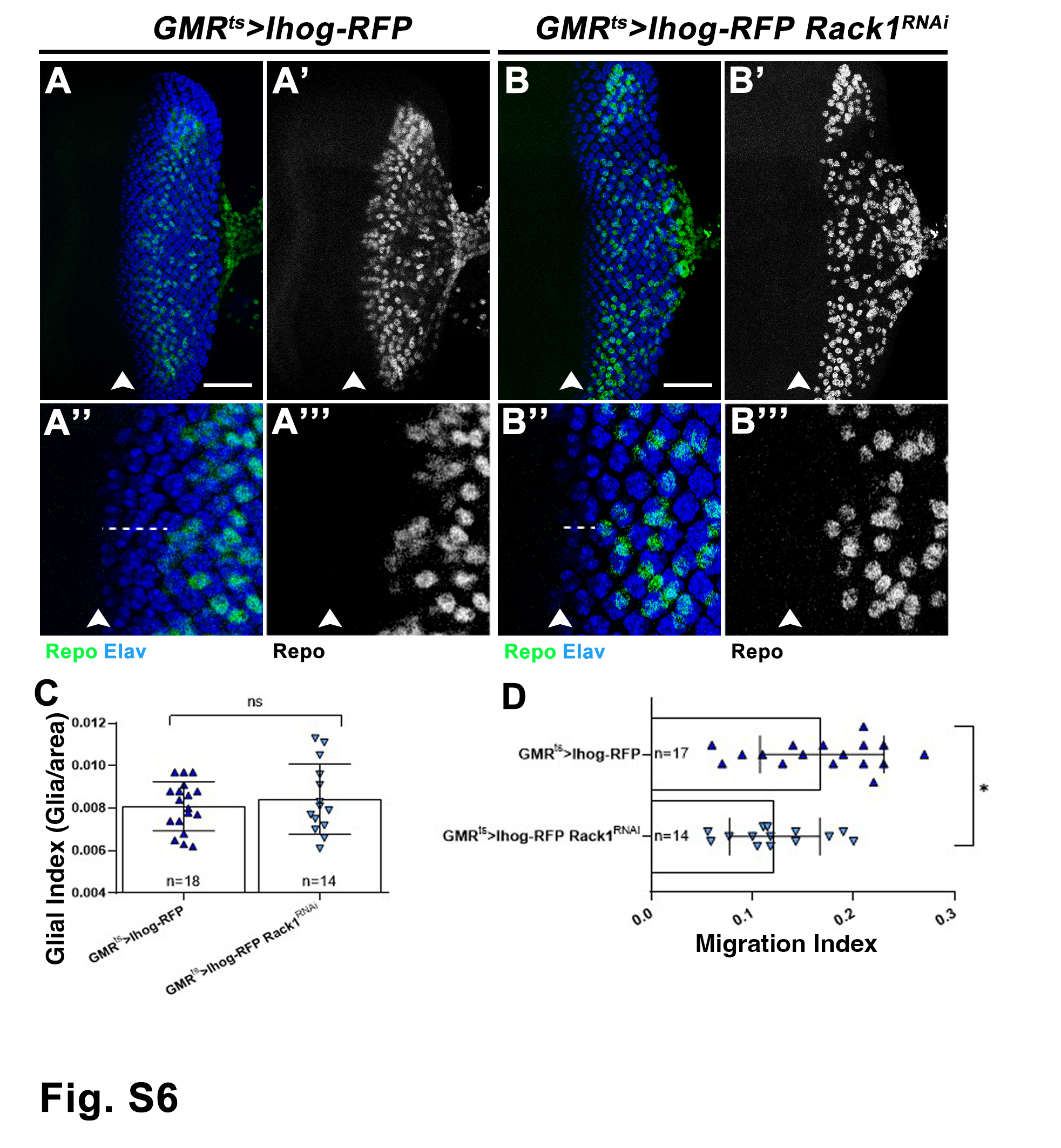

### Fig S7

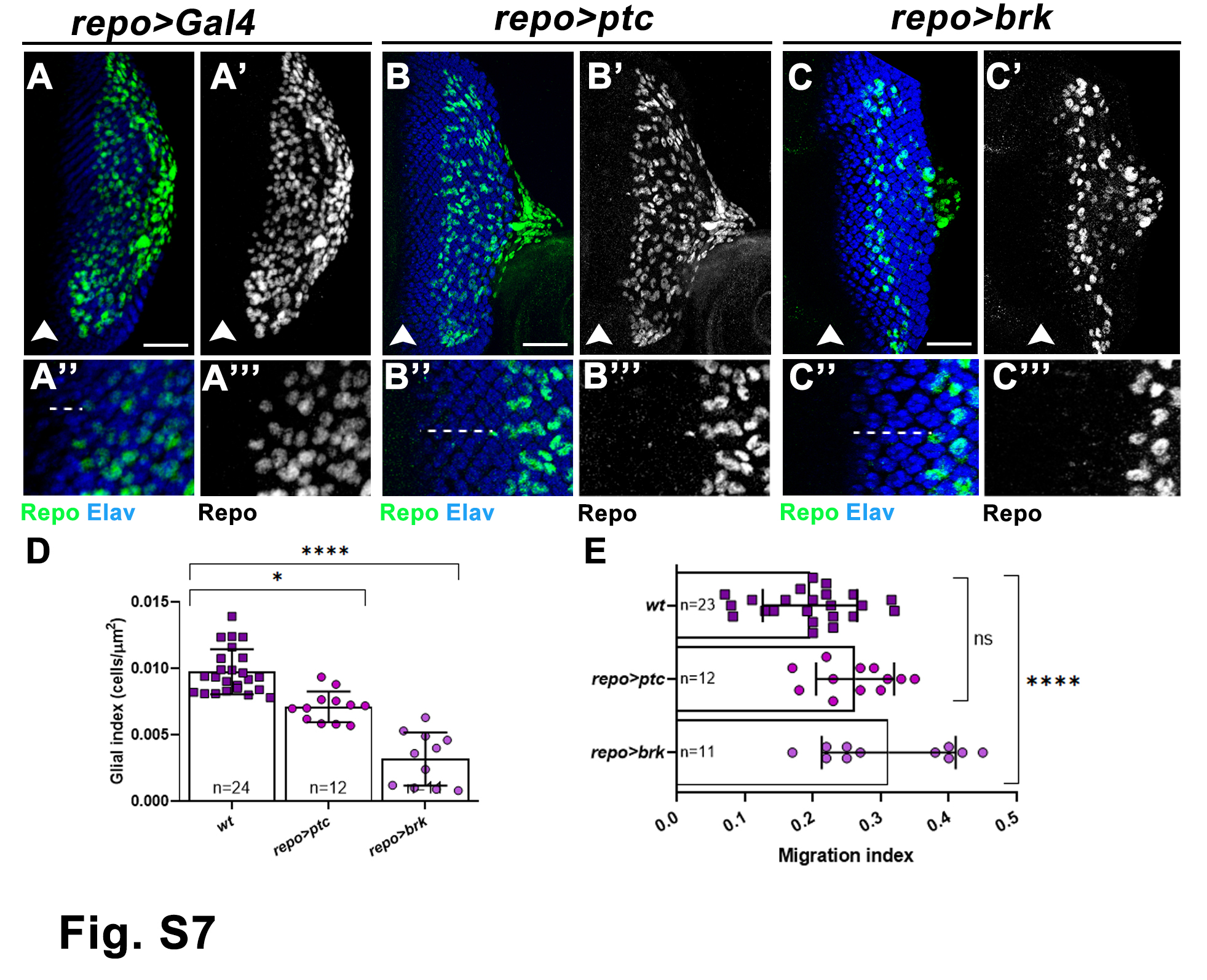

### Fig S8

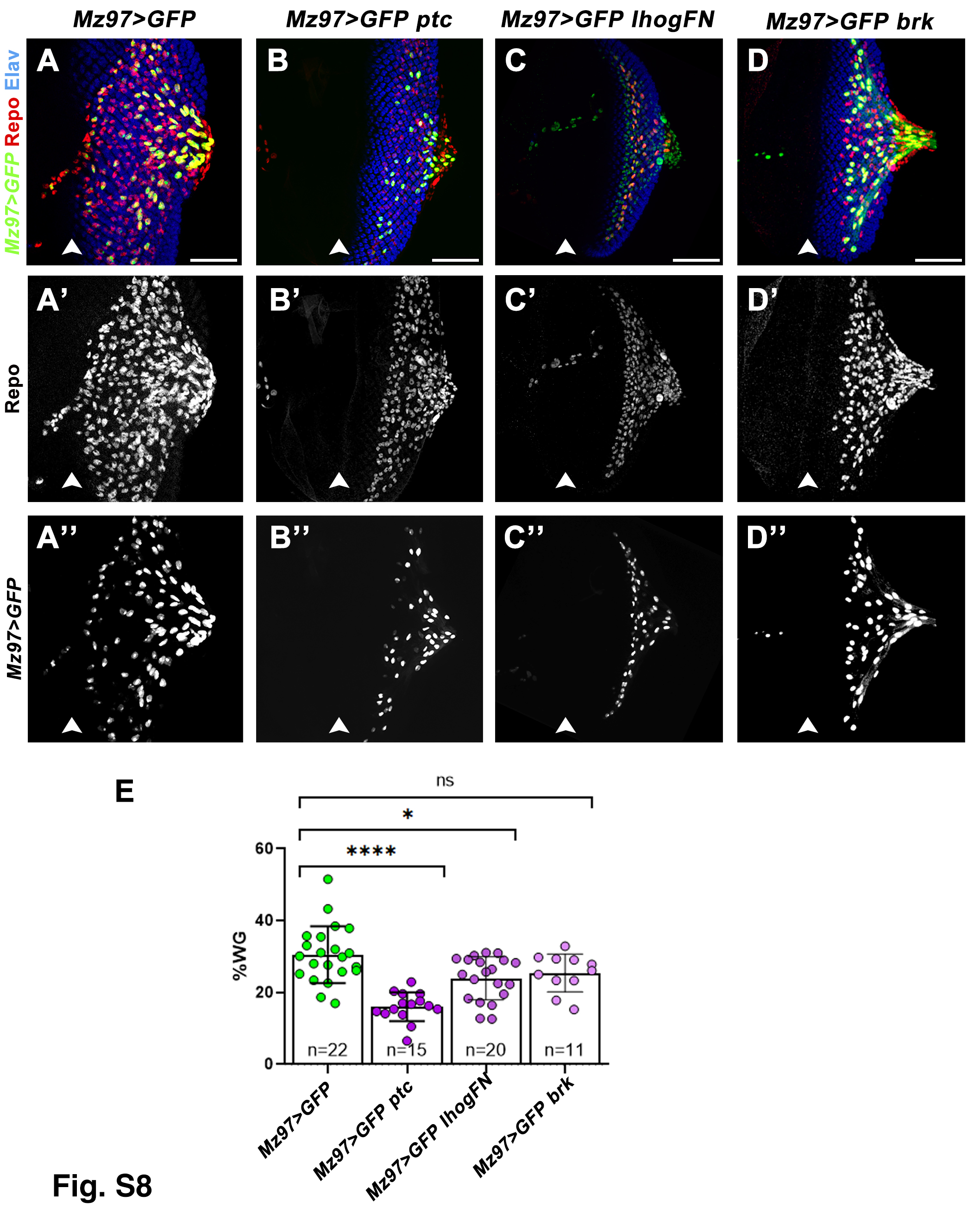

### Fig S9

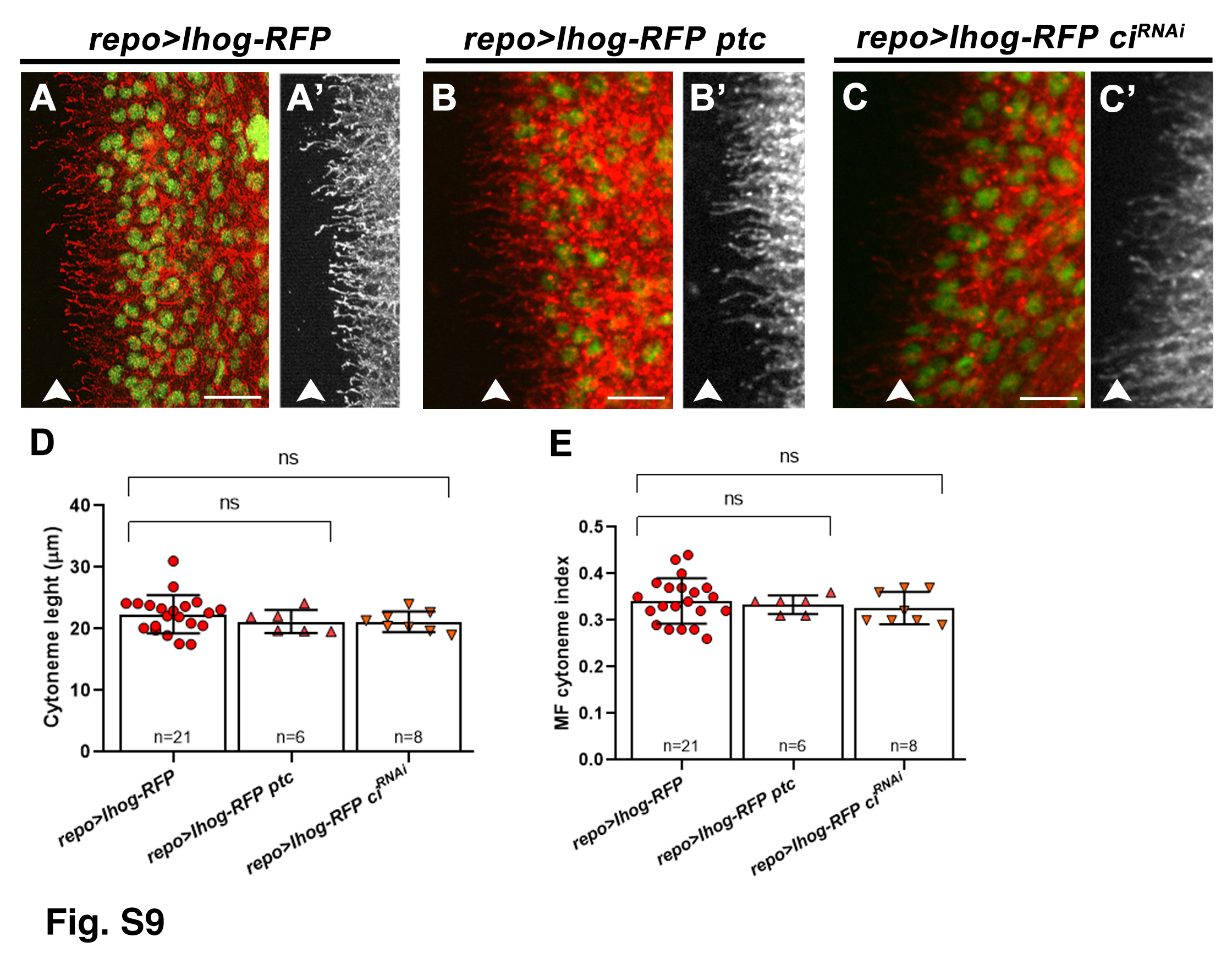
